# Characterization of a novel heat tolerance trait and subsequent haplotype block-based analysis to identify causal regions in Dutch Holstein cattle

**DOI:** 10.1101/2025.05.02.651836

**Authors:** T. Pook, M. L. van Pelt, J. Vandenplas, I. Adriaens, L. Zetouni, C. Orrett, Y. de Haas, C. Kamphuis, B. Gredler-Grandl

**Affiliations:** Wageningen University and Research, Animal Breeding and Genomics, P.O. Box 338, 6700AH Wageningen, the Netherlands; Cooperation CRV, Animal Evaluation Unit, P.O. Box 454, 6800 AL Arnhem, the Netherlands; CRV BV, Global Genetics R&D, P.O. Box 454, 6800 AL Arnhem, the Netherlands; KU Leuven, Animal and Human Health Engineering Division, Kleinhoefstraat 4, 2440 Geel, Belgium

**Author notes:** Corresponding author E-mail addresses: TP.

**Keywords:** heat tolerance, heat stress, haplotype block, GWAS

## Abstract

Heat stress is a major environmental challenge affecting dairy cattle, leading to behavioral changes, production losses, and welfare concerns. As heat stress events intensify and become more frequent due to climate change, identifying heat tolerant animals is crucial for sustainable dairy production. This study develops a pipeline to quantify the population-wise impact of heat stress on a dairy cattle population and subsequently defines individual-based heat tolerance traits. Data from 677,318 Dutch Holstein cows, including 15.6 million mid-infrared spectra and 762 million records from automated milking systems, were analyzed. An iterative approach using kernel regression was employed to estimate the population-wise effects of heat stress. Results indicate that fat and protein percentages decrease approximately linearly with increasing temperature humidity index (THI) with an absolute reduction of 0.3% from THI = 30 to THI = 70. In contrast, milk yield remains stable until a THI of 60, after which production losses increase quadratically reaching 5.0% at a THI of 75. The phenotype of an animal is subsequently defined as the slope in a linear regression model of the residuals of the population-wise models against THI for milk yield and concentration of fat, protein, lactose, and specific fatty acids. Compared to reaction-norm models, individual records per cow are combined before model fitting, thereby reducing computation times and allowing more flexibility in the design of the model. Heritabilities for heat tolerance traits ranged from 0.05 to 0.12, and genetic variances indicate substantial potential for breeding as an improvement of the population by one genetic standard deviation would already offset 69% of the losses in fat percentage, 65% in protein percentage, and 11% in milk yield. Heat tolerance based on milk yield showed favorable correlations with most commercial traits, whereas heat tolerance based on fat and protein percentage showed negative correlations to health and resilience. A genome-wide association study using both SNPs and haplotype blocks from the software HaploBlocker identified potential QTLs across the genome, with particularly strong signals on BTA5, 14, and 20. These findings support the potential of breeding for heat tolerance but highlight the need for complementary management strategies to mitigate heat stress impacts.

**Interpretive summary:** This study introduced a novel, computationally efficient method to quantify the impact of heat stress in dairy cattle and define novel heat tolerance traits based on milk production data from automated milking systems. Our results indicate quadratically increasing losses in milk yield with increasing heat load. The identified heat tolerance traits show substantial genetic variance, moderate heritabilities, and favorable correlation to key production traits. These findings highlight the potential for incorporating heat tolerance into dairy breeding goals to mitigate climate change impacts, improve animal welfare, and enhance sustainable milk production.

## Introduction

Heat stress is an important environmental challenge that affects dairy cattle and has profound physiological impacts resulting in behavioral changes, productive losses, and animal welfare issues (Marino and Allen, 2017; Herbut et al., 2021). As global temperatures continue to rise due to climate change (Cheng et al., 2022), the frequency and intensity of heat stress events are projected to increase, exacerbating these impacts (Pörtner et al., 2022). This, in turn, causes economic losses not only in dairy cattle production but also in other livestock species worldwide (St-Pierre et al., 2003; Ross et al., 2015; McManus et al., 2020).

Various measures to combat heat stress have been developed across different domains of the animal production sectors. This includes adapting nutrition strategies to optimize diets and enhance energy availability to reduce metabolic heat production (Ríus, 2019), as well as modifying housing and management systems to implement cooling strategies (Johnson, 1987; D’Emilio et al., 2017), such as the use of fans, sprinklers, or shade structures.

In this manuscript, we will focus on the use of breeding and genetic improvement as an additional measure to combat heat stress. Genetic differences in the handling of heat stress between breeds are well established (Copley et al., 2024) and are utilized in crossbreeding when introducing genetic material to low-production environments while maintaining local adaptation (Jordan, 2003; Michael et al., 2021). However, relatively little attention in applied breeding is given to within-population differences. Australia is the only country that explicitly includes heat tolerance in its selection index (Nguyen et al., 2016), while the Netherlands incorporates a general resilience trait in the dairy cattle breeding goal (Poppe et al., 2022). Although genetic improvement has less of a short-term impact, it remains an important consideration for a long-term sustainable breeding goal (Ravagnolo and Misztal, 2000; Garner et al., 2016).

Heat stress in this context is defined as an environmentally challenging condition. For the quantification of this condition, there are various approaches with the easiest one being the use of weather data such as the daily peak temperature or humidity (National Research Council, 1971). To quantify the challenge level of an animal, potential indicator traits such as increased rectal temperature and respiratory rate (Li et al., 2020; Yan et al., 2021) can be used. For the purpose of breeding, the goal is subsequently to define a trait to describe how well the animal is responding to heat stress, aiming either for heat tolerance or heat resilience (Misztal et al., 2024). In short, heat tolerance describes the ability of an animal to maintain its production levels during heat stress, while heat resilience describes the ability of an animal to quickly return to its production level after heat stress.

To quantify the challenge level of an animal, potential indicator traits such as increased rectal temperature and respiratory rate (Li et al., 2020; Yan et al., 2021) can be used. As the large-scale recording of rectal temperature and respiratory rate is impractical in the commercial setting due to high cost, more and more attention is given to the use of routinely collected traits such as milk yield to measure the impact of heat stress across lactations, weather conditions, and production environments to model genotype by environment interactions (GxE) (Kipp et al., 2021; Vinet et al., 2023). Currently, two main approaches are used in animal breeding to model GxE interactions, and more specifically heat stress: First, multi-trait models are applied, where traits recorded in different environments are modelled as being genetically correlated traits (Falconer, 1952). Practical analysis suggests measurable benefits in this approach when genetic correlations fall below 0.8 (Copley, 2024). Second, reaction-norm models are employed to estimate interaction effects between traits and environmental conditions (Calus and Veerkamp, 2003; Kolmodin et al., 2002). Weather conditions have been shown to strongly affect fertility traits (Ojo et al., 2024), however, correlations between milk yield traits in different weather conditions are usually high, failing to separate heat tolerance from overall production levels (Aguilar et al., 2010).

On a technical level, a reaction-norm model is a linear mixed model (Henderson, 1975) with this or similar structure (Su et al., 2006):

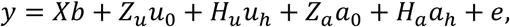

where X, Z, and H are incidence matrices for the fixed effects (b), breeding values for the intercept (𝑎_0_) and slope (𝑎_ℎ_) of the reaction to environmental conditions, and non-genetic components 𝑢_0_ & 𝑢_ℎ_ that are modelled as random effects. Here, intercept and slope components are usually modelled as correlated in a multivariate normal distribution with respective correlations 𝜎_𝑢0,𝑢ℎ_ and 𝜎_𝑎0,𝑎ℎ_. The interested reader is referred to Carabano et al. (2017) for an extended overview of reaction-norm models.

Reaction-norm models are powerful tools capable of accurately estimating GxE effects, but they require large datasets. Since reaction-norm models inherently include the heat tolerance (slope) as a random effect, they require prior estimation or assumption of variance components and process each record of an animal separately. This makes them computationally expensive and limits their applicability in downstream analyses such as genome-wide association studies (Holland and Piepho, 2024).

The aim of this study is to develop a pipeline for the estimation of the population-wise and individual-based impact of heat stress on milk traits in dairy cattle. A further aim for this pipeline is to be both computationally efficient and able to include non-linearity and interaction between different parameters. Based on this, we are defining novel heat tolerance traits and the potential impact of breeding through genetic improvement is critically assessed by estimating variance components and identification of causal regions by use of a genome-wide association study and a haplotype block-based analysis.

## Materials and Methods

In the following, we provide a pipeline to estimate population-wise effects of heat stress, including pre-processing steps from common input data formats. Based on this, we define new heat tolerance traits and subsequently assess them in detail by estimating variance components, performing a genome-wide association study (GWAS), and a haplotype block-based analysis. A schematic overview of the pipeline is provided in Figure 1.

**Figure 1:**
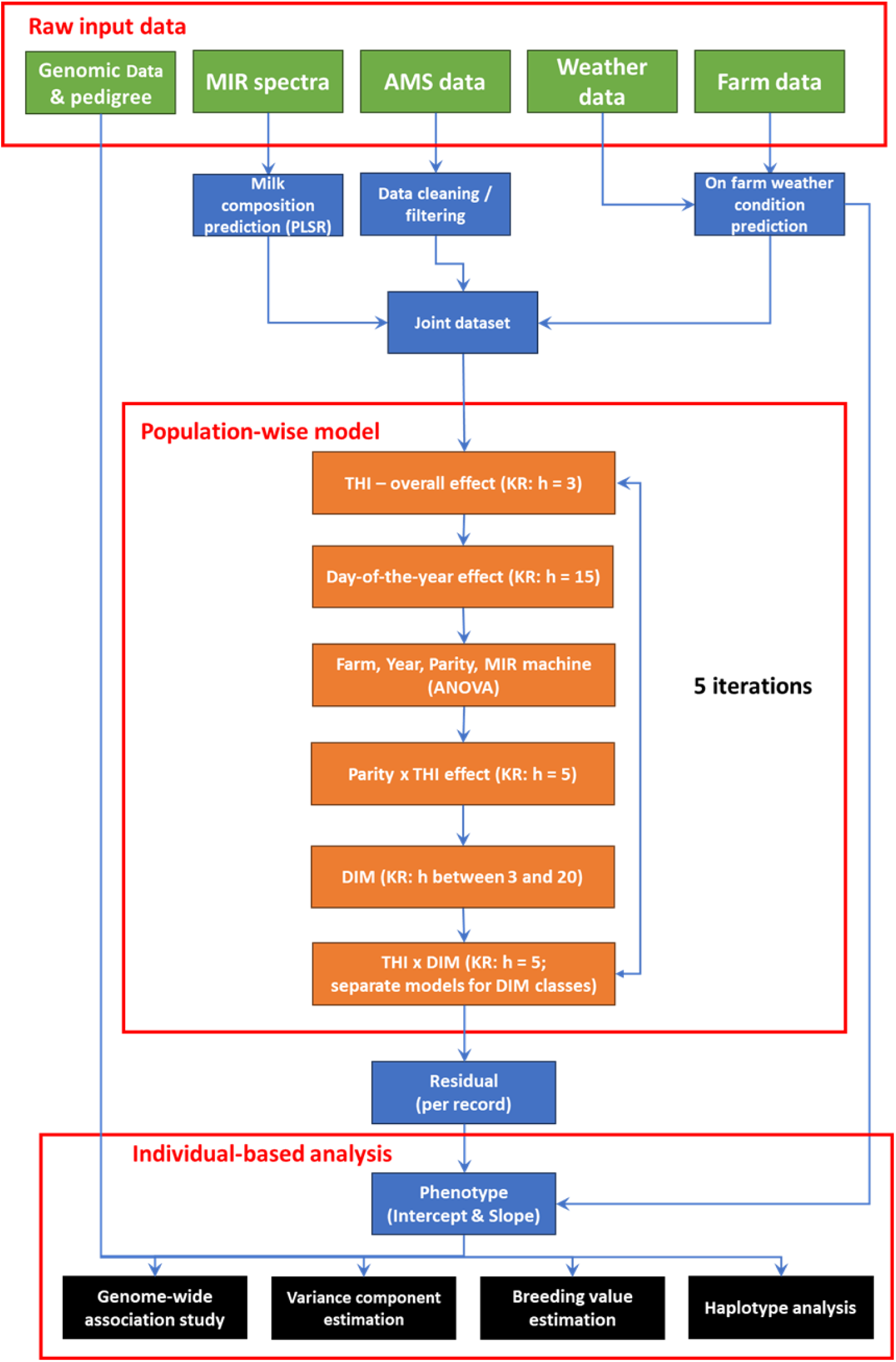
Schematic overview of the processing pipeline for the estimation of population-wise effects of heat stress and subsequent individual-based analysis.

### Materials

Data from the Dutch dairy cattle breeding company CRV (Arnhem, the Netherlands, www.crv4all.nl) was used, including 677,318 animals from 1,478 farms spread across the Netherlands and covering a period from 2013 to 2021. Of these animals, 348,133 were genotyped using different genotyping arrays, primarily using the Illumina EuroG MD chip (https://www.eurogenomics.com/eurog-md-chip.html), which was subsequently imputed to obtain a joint panel of 76,438 SNPs. Furthermore, 5,322,943 animals were included in the pedigree, tracing back up to 30 generations till 1910. Individual Mid-infrared (MIR) data were generated through routine milk recording procedures (ICAR, 2023) by QLIP (Zutphen, the Netherlands, www.qlip.com) with spectra per cow being generated monthly, resulting in a total of 15,596,136 MIR spectra. Although no milk composition data were available for this dataset, estimates for fat percentage (F%), protein percentage (P%), lactate, and individual fatty acids were generated using a smaller in-house dataset including 1,740 MIR spectra with the respective information, using partial least squares regression (PLSR, (Soyeurt et al., 2006)). Details on the estimation procedure are provided in Supplementary File S1 and Supplementary Figures S1 and S2. Furthermore, milk yield data were collected using automated milking systems (AMS) with 762,173,326 records from individual milkings across 934 farms.

For all subsequent analyses, the dataset was reduced to a panel of animals for which data from all data sources were simultaneously available in sufficient quantity (at least five MIR spectra per animal), resulting in a panel of 5,929,221 MIR spectra from 346,248 animals across 772 farms (Table 1).

**Table 1:** Overview of the dataset used.

| Year | Number of spectra | Number of animals | Number of farms | Records (THI $> 60$) | Records (THI $> 70$) |
| --- | --- | --- | --- | --- | --- |
| 2013 | 679.485 | 210.399 | 738 | 138.839 | 7.111 |
| 2014 | 700.179 | 220.893 | 749 | 145.835 | 4.204 |
| 2015 | 743.947 | 235.003 | 759 | 124.920 | 3.206 |
| 2016 | 817.012 | 255.491 | 762 | 216.088 | 10.514 |
| 2017 | 719.829 | 255.491 | 764 | 180.272 | 1.260 |
| 2018 | 717.618 | 245.471 | 765 | 230.861 | 11.114 |
| 2019 | 708.815 | 239.885 | 758 | 160.580 | 14.473 |
| 2020 | 571.811 | 248.317 | 752 | 111.043 | 8.588 |
| 2021 | 270.524 | 247.990 | 745 | 39.599 | 393 |

Meteorological data from the Copernicus Climate Change Service (C3S, (Copernicus Climate Change Service, 2024)) was used, gridded at a 0.1-degree resolution for longitude and latitude, with information on temperature, precipitation, wind, pressure, and humidity based on satellite observations, ground-based weather stations, and climate models. For reference, 0.1 x 0.1 degrees corresponds to an approximately 11 x 7 km grid in the Netherlands, providing a much finer grid than the use of weather stations from the Koninklijk Nederlands Meteorologisch Instituut, as used in Ojo et al. (2024), to derive the temperature humidity index (THI, (National Research Council, 1971)). Weather data for each farm were subsequently approximated using the weather conditions of the closest data point, based on the global positioning system (GPS) coordinates of the farms (Figure 2). As the Netherlands is very flat and the used grid is narrow these values should be highly representative for each specific farm. It should be taken into account, that for more hilly regions, a more sophisticated approach to combine data from multiple weather stations in the area (Gote et al., 2024) or weather recording on farm might be necessary. Of all records, 22.7% were in conditions above THI = 60, while only 1.03% were above THI = 70. In 2017, there were essentially no extreme heat waves with conditions above THI = 70. Whereas MIR spectra were approximately equally distributed across the years from 2013 to 2020, fewer records were available in the summer of 2021, resulting in just 15.9% of records being above THI = 60 in 2021.

**Figure 2:**
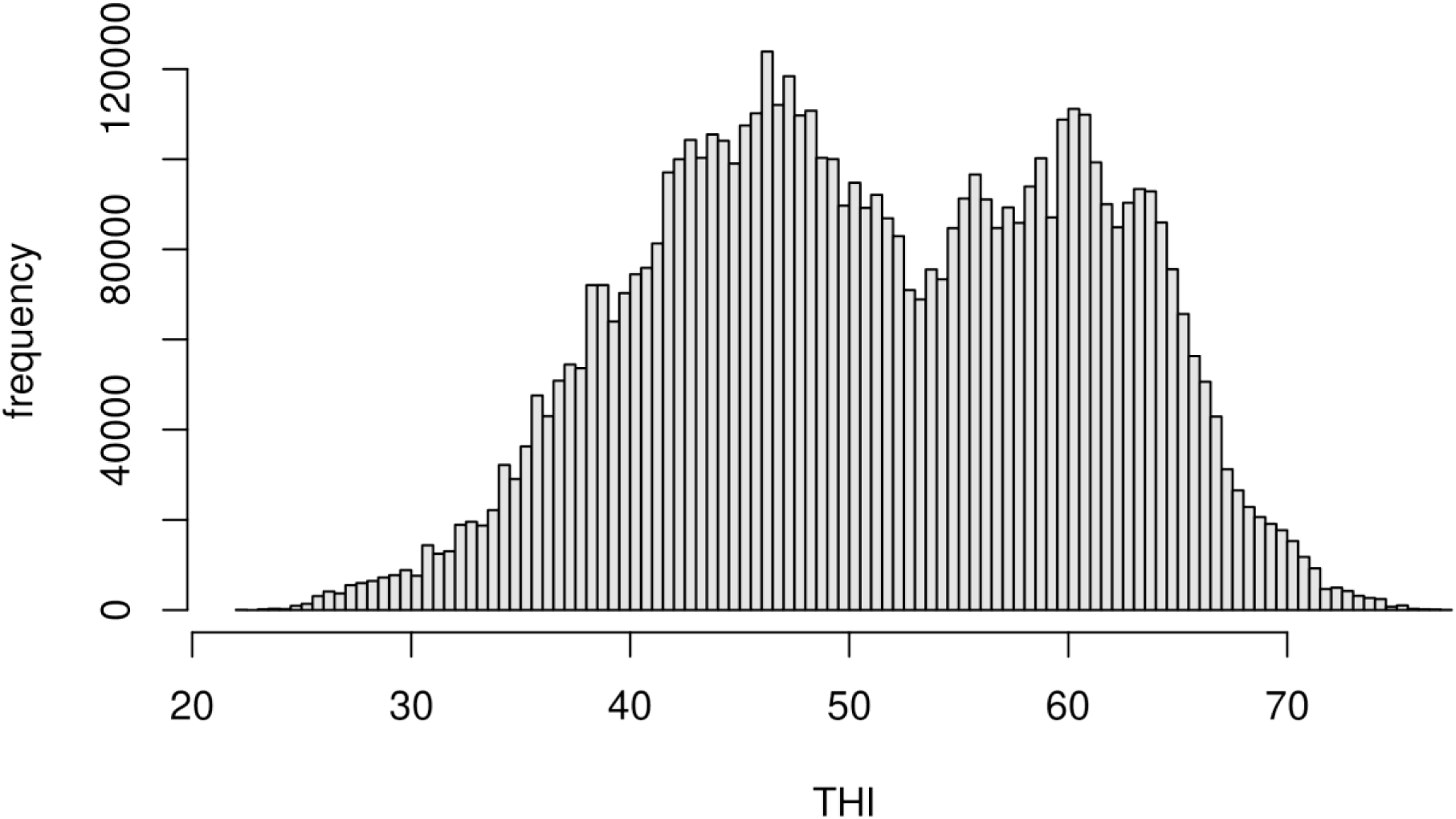
Distribution of weather conditions based on THI of the last three days (including the same day) for the used MIR records based on the Copernicus Climate Change Service (C3S, (Copernicus Climate Change Service, 2024)) and GPS coordinates of farms.

### Pre-Processing of data

Before conducting the main analyses, we pre-processed the data to ensure consistency and reliability. This involved calculating THI values, filtering milk yield data, and applying statistical smoothing techniques.

THI was calculated from the weather data using the average daily temperature in Celsius (T) and relative humidity (RH) (National Research Council, 1971):

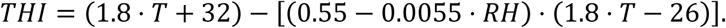

In all subsequent models, the average THI from the last three days (including the current day) was used in accordance with Mattalia et al. (2023). When referring to THI in the rest of the manuscript, the average THI from the last three days is meant.

Data from AMS were aggregated to calculate daily milk yield per animal. Basic data cleaning protocols, similar to those used at CRV for routine genetic evaluations, were employed to avoid double-counting of records, among other issues. Additionally, strict filters were applied to focus on “normal” lactations, only considering lactations with an overall length between 200 and 450 days and milk yield greater than 0 and less than 60 liters for at least 95% of the days in lactation. If milk yield changed by more than 10 liters from one day to the next, records from one week before and after these cases were excluded, as no reliable health data were available to correct for such outliers. Additionally, the first 10 days and the last 5% of lactation days were excluded to avoid issues with the fit of the lactation curve.

Based on the remaining data, the lactation curve *l* for each individual cow was estimated using a kernel regression-based approach, using a Nadaraya-Watson estimator (NW) with a Gaussian kernel and a bandwidth of 25 (Nadaraya, 1964), based on the interquartile range:

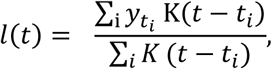

With 𝑡_𝑖_ being the number of days in milk (DIM) and 𝑦_𝑡𝑖_ being the milk yield on the given day. The relative performance of each cow, compared to its lactation curve, was derived as the ratio of the current and expected performance. To minimize the impact of non-systematic daily fluctuations, current performance was smoothed using a Nadaraya-Watson estimator with a reduced bandwidth of 3. Milk data on previous days were excluded, as heat stress on a given day does not impact yield on the previous day:

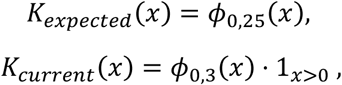

where 𝜙_𝜇,𝜎_ is the density of a Gaussian distribution with mean 𝜇 and standard deviation 𝜎.

Instead of using the aggregated fat or protein yield in kilogram as traits, we consider the relative yield per individual, compared to its own expectation by scaling F% and P% using previously calculated ratio l(t), which will subsequently be referred to as relative yield (RY_F & RY_P). No smoothing was applied to F% and P% as only monthly measurements were available.

### Population-wise effects of heat stress

For the estimation of population-wise effects of heat stress, a combination of parametric and non-parametric modelling approaches was used that was performed iteratively by using the residuals of the previous step as the response variable in the next step (Figure 1).

Firstly, the overall effect of THI was estimated using a NW estimator with a bandwidth of 3. Subsequently, a day-of-the-year effect was estimated using a NW estimator with a bandwidth of 15. Next, fixed (class) effects for farm, year, parity, and machine (used to generate the MIR spectra) were estimated using a linear regression model. Parities 4 and higher were considered as one class. Next, THI by parity effects were estimated using four separate NW estimators, in which only reports from parity 1, 2, 3, or ≥4 respectively, were used. As fewer records compared to the overall THI effect were available, a bandwidth of 5 was used. Next, an effect for DIM was fitted again using a NW estimator. For DIM ≤ 350 a bandwidth of 3 was used, however, as fewer observations were available later in lactation and only limited differences in this phase are expected, a bandwidth of 20 was used for DIM > 350. Lastly, an interaction effect between DIM and THI was fitted by assigning observations to separate DIM classes, for which separate THI effects with a bandwidth of 5 were estimated: 30 or fewer days, 31-100 days, 101-200 days, 201-300 days, 301 or more days.

This iterative procedure was repeated five times. Subsequently, THI x DIM and THI x parity effects were centered around zero by adapting the overall THI effect by the weighted mean of the effects for the different classes of DIM and parity (based on frequency) and the required change added to the overall THI effect.

The aforementioned bandwidths were chosen based on visual inspection. Larger (future) datasets may allow for reduced bandwidths, whereas smaller datasets might require increased bandwidth to reduce prediction variance (Hassanpour et al., 2023). Initially, models included an adaptive choice of bandwidth to use larger bandwidths in areas with fewer observations, however, this was later dropped to simplify models and stabilize prediction in areas with a lower number of observations (Brockmann et al., 1993). No multivariate NW estimators (Nadaraya, 1964) were used as this would have required an increase of bandwidths to obtain similar robustness of fit and the number of observations in extreme THI values was limited to begin with.

### Individual-based heat tolerance

After estimating population-wise heat stress effects, we defined heat tolerance traits by modeling residual variations at the individual level. The individual phenotype of an animal for the heat tolerance trait is defined based on its relative performance compared to the expected performance estimated using the previously derived population-wise model. To do this, the finally obtained residuals of the animal from the population-wise models are regressed against THI in a linear regression model with the obtained slope of the regression model being the phenotype for the heat tolerance trait. THI values were adjusted by 50 so that the intercept of the regression model represents the performance of an animal at THI = 50, which should represent its performance at thermo-neutral conditions with the average THI of 51.1 across all samples and 47.8% of all records being generated at THI > 50. An exemplary visualization of a single cow is given in Figure 3. Note that intercept and slope are explicitly not the result of a random regression model in which population-wise effects and intercept and slope are jointly estimated as in a reaction-norm model (Su et al., 2006).

**Figure 3:**
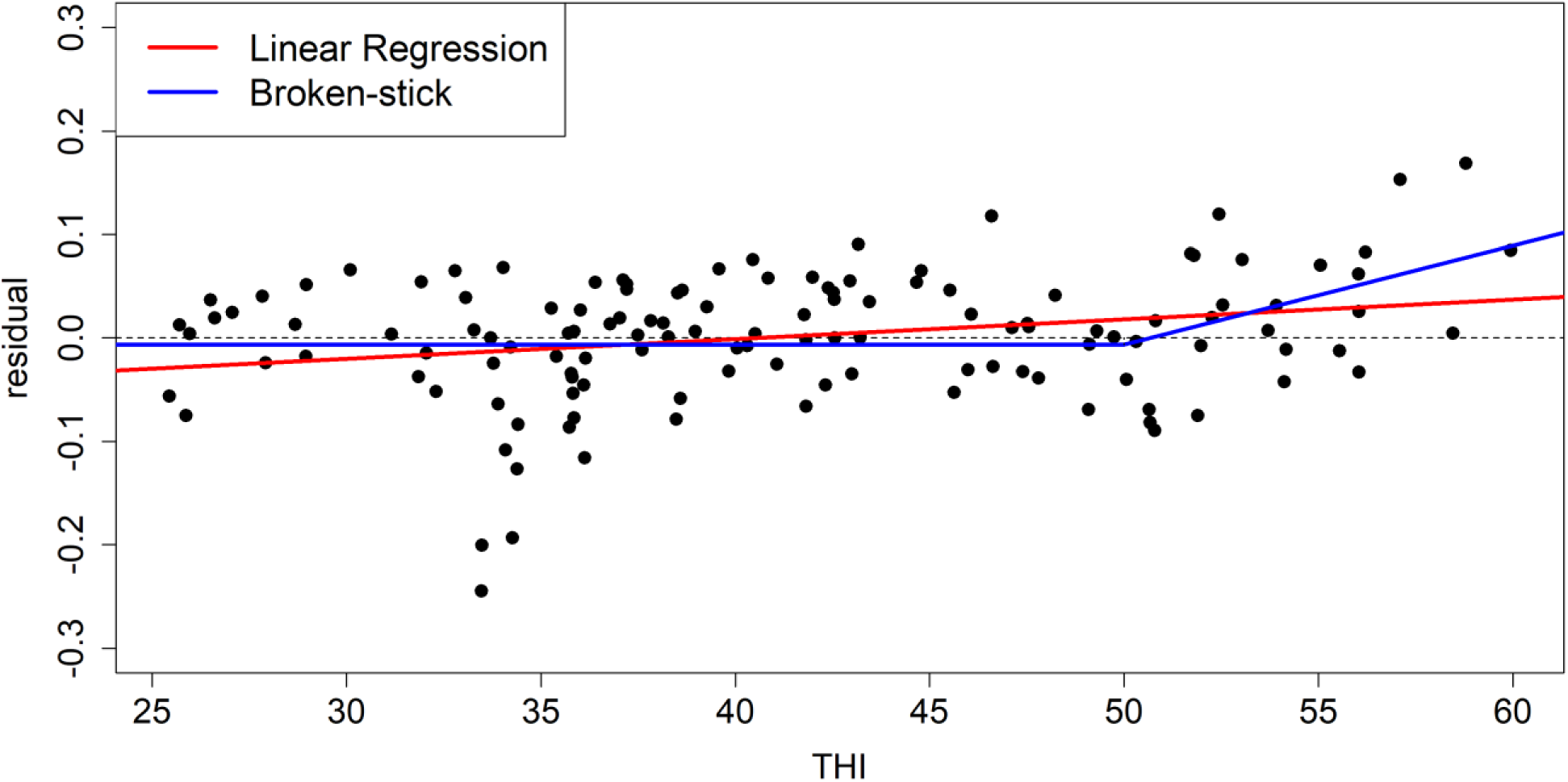
Exemplary visualization of the calculation of the phenotype for heat tolerance based on milk yield. Dots represent individual observations for the same cow.

To avoid a strong impact from records in cold weather conditions, a second version of the trait was calculated in which observations below a minimum THI value were set to that value, mimicking a broken-stick model in a reaction-norm model. Different THI values were considered as thresholds, with THI = 50 used as a baseline model (alternatives: THI = 40, THI = 45, THI = 55). As there were substantially more records for milk yield, a higher threshold (THI = 60) was considered here as well. Only cows with a minimum of 5 records (or 50 in the case of milk yield) were given a phenotype to ensure reasonable reliability in the estimation of the regression coefficients. Furthermore, alternative THI thresholds of a minimum of 7, 10, 12, 15, 20, and 25 records were considered.

### Variance component estimation

To assess the potential of using the previously defined heat tolerance traits for breeding, genetic variance components of traits were estimated using univariate models including a fixed effect for farm (class effect) and performance in thermoneutral conditions (intercept) as a covariance using ASRemlR (Butler et al., 2009). To increase model robustness, phenotypes with an absolute value above the 99% quantile were set to the 99% quantile with appropriate signs to reduce the impact of extreme phenotypes. Variance components were estimated based on pedigree data, including all phenotyped animals traced back for three generations. Genetic correlations between heat tolerance traits were subsequently estimated pairwise in bivariate models based on pedigree using ASRemlR (Butler et al., 2009). Genetic correlations to other important breeding goal traits were estimated based on multiple across-country evaluation (MACE) correlations (Schaeffer, 1994) following the approach described in Poppe et al. (2022)

### Genome-wide association study

In addition to the 76,438 SNPs, haplotype blocks derived using the software HaploBlocker (Pook et al., 2019) were considered in a genome-wide association study. In short, a haplotype block in HaploBlocker is a specific sequence of alleles and only those individuals that carry this sequence in one of its two haplotypes (phased genotype) is carrying the given haplotype block. This effectively screens the population for cases of group-wise identity by decent (Donnelly, 1983), allowing haplotype blocks to overlap with potentially unique start and end points of each haplotype block, leading to longer block structures than identified with conventional haplotype block detection approaches (Barrett et al., 2005).

For this, genotypes were first phased using Beagle v5.4 (Browning et al., 2021) (ne = 1,000 (Pook et al., 2020)), and subsequently, haplotype blocks were derived in HaploBlocker using window_size = 10 and min_majorblock = 1,000. Btau 4.0 (Bovine Genome Sequencing and Analysis Consortium et al., 2009) was used as the reference genome for both phasing and haplotype block calculation.

The genome-wide association studies for our newly proposed heat tolerance traits were performed using the R-package statgenGWAS (van Rossum et al., 2020) including a fixed effect for both farm and performance in thermoneutral conditions (intercept). To reduce computing time, the data was split into six subsets, and p-values from the subset analyses were combined using the weighted Z-score method (Willer et al., 2010).

To aid results from the GWAS, phenotypic differences between carrier and non-carriers of specific haplotype blocks were estimated using a linear regression of the observed phenotypes against haplotype block count (0, 1, 2). Specifically, direct estimation of haplotype blocks or as the sum of the effects of the included SNPs using a mixed model was avoided (Meuwissen et al., 2001), due to potential biases in estimation caused by LD and higher impact of shrinkage on rare variants.

## Results

### Population-wise models

Results from the population-wise models indicate an essentially linear decrease in both F% (Figure 4.A) and P% (Figure 5.A) with increasing THI. The effect of THI is approximately 20% smaller for first parity cows with a loss of 0.29% and 0.20% for F% and P% from THI = 30 to THI = 70, respectively, compared to later parities with average losses of 0.35% and 0.25%. Note here that overall concentration levels for first parity cows are slightly lower. Furthermore, the season effect for both F% and P% in June and July is negative, resulting in further reduction of F% and P% (-0.1% / -0.06%) with substantially increased P% in autumn (up to +0.08%), while F% for autumn until the end of winter is quite constant at +0.03% (Figures 4.C and 5.C). The effect of DIM on F% and P% is substantial, with expected F% and P% around day 50 of lactation being almost 1% lower than for the very beginning and end of lactation (Figures 4.D and 5.D). In comparison, the absolute size of the interaction effect of THI and DIM is relatively low. Nonetheless, models indicate a lower impact of heat conditions on P% in early lactation as the positive effect is complementary to the overall negative effect of THI on P%. Therefore, all these effects need to be interpreted as residual effects. For example, a cow is expected to have 0.061% reduced P% at THI = 60 (Figure 5.A), however, if we additionally know that the cow is in the first thirty days of lactation (+ 0.043 at THI = 60, Figure 5.B), the expected overall impact of weather conditions on that cow on the given day is the sum of both effects (-0.017%; excluding the seasonal effect).

**Figure 4:**
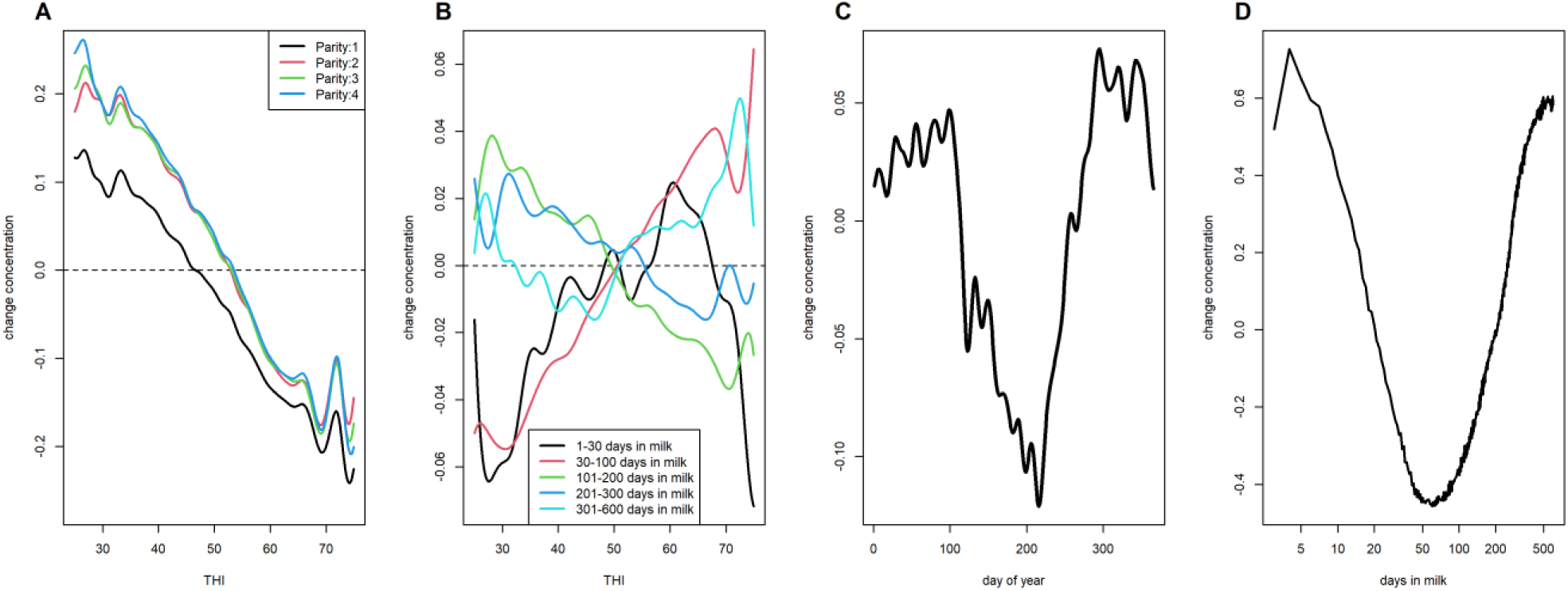
Estimated effects of THI x parity (A), THI x DIM (B), season (C), and DIM (D) on fat percentage in milk.

**Figure 5:**
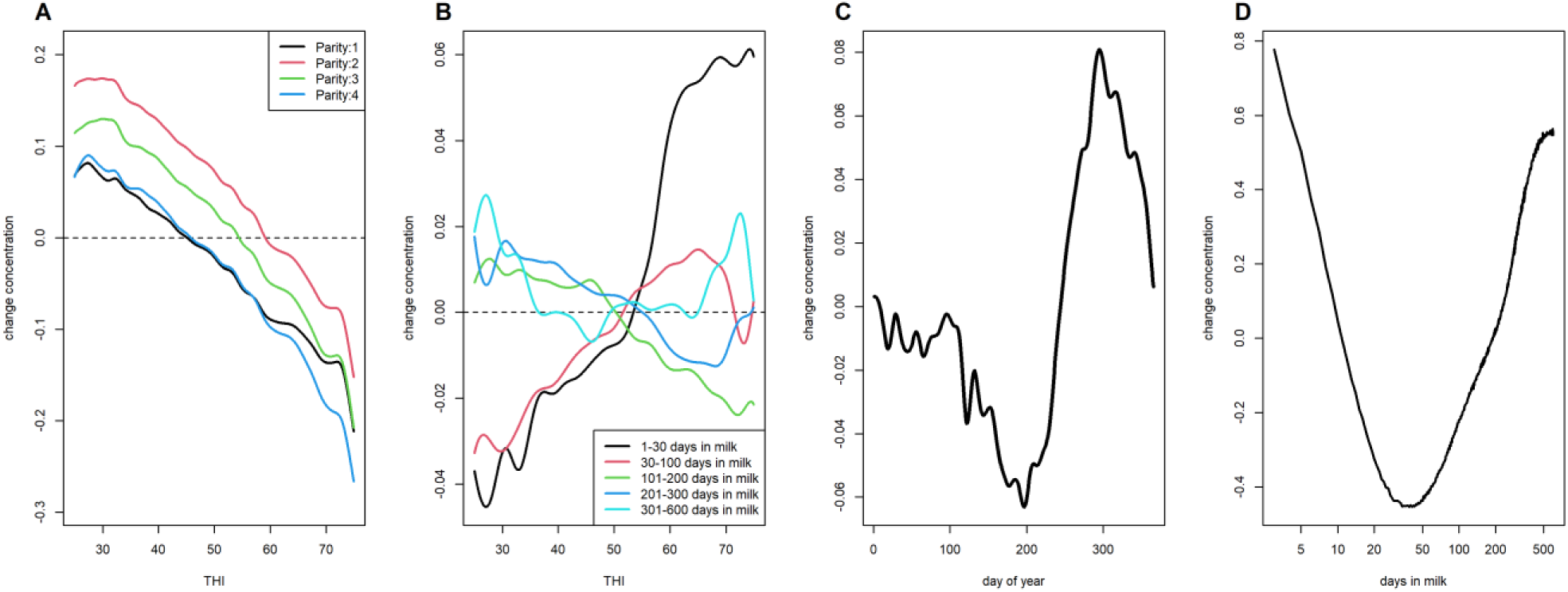
Estimated effects of THI x parity (A), THI x DIM (B), season (C), and DIM (D) on protein percentage in milk.

The interested reader is referred to Supplementary Figures S3 to S14 with results on lactose and the individual fatty acids. As an example, consider the often with heat stress associated fatty acids C-18:1 cis9 (Hammami et al., 2015), which is shown to substantially increase in concentration for cows in early lactation (< 30 days, Figure 6.D) as well as in first parity cows overall. Furthermore, the estimated THI x DIM interaction effect suggests an even stronger increase under heat conditions (THI > 65) in cows within the first 100 days of lactation (Figure 6.B).

**Figure 6:**
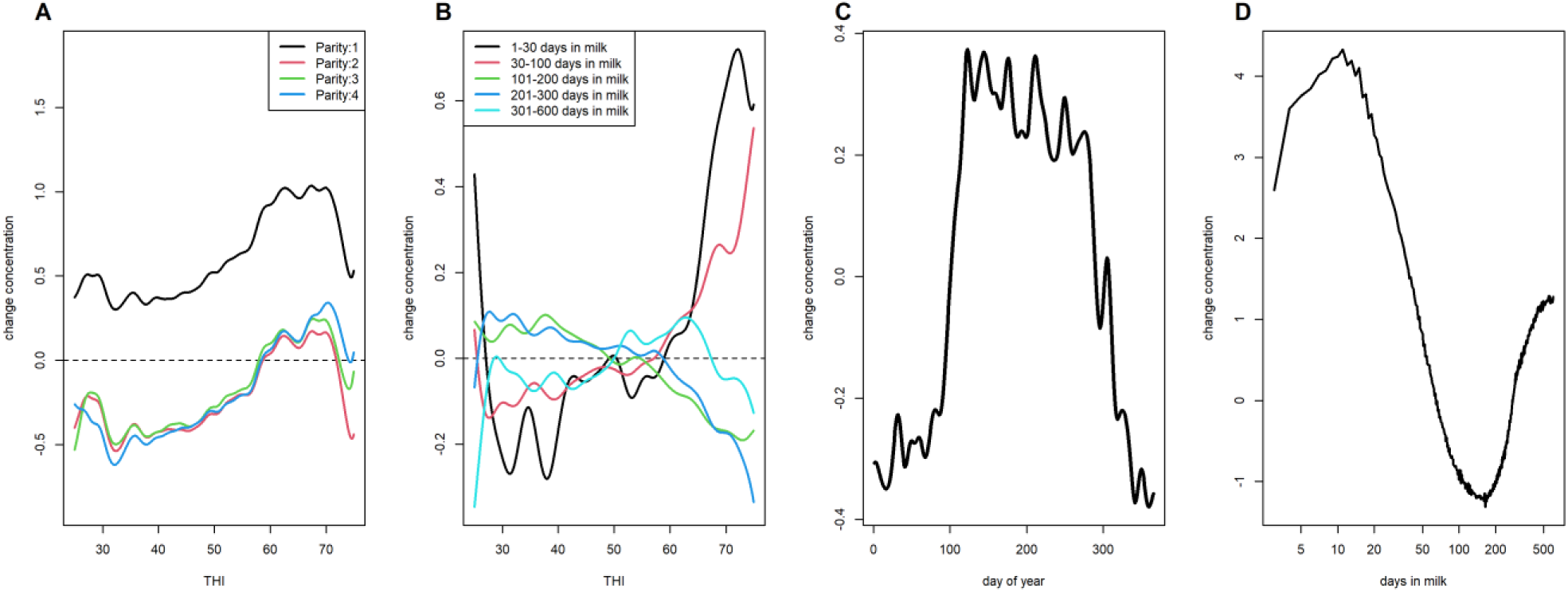
Estimated effects of THI x parity (A), THI x DIM (B), season (C), and DIM (D) on C-18:1 cis9 concentration in milk.

Results reported in this study were obtained using a joint model including data from all parities, however, fitting separate models for data from each parity resulted in overall similar estimates for all effects (Supplementary Figures S15 and S16). Noticeably, effect estimates for the THI x DIM effect for the individual parities showed high variance in the fit, indicating the increased power by the use of a larger joint dataset (Supplementary Figure S17).

Models for milk yield suggest that yield is increasing up to THI 55 (Figure 7.A). Therefore, partially compensating for reduced F% and P% at moderately increasing THI. However, as THI increases milk yield drops by 0.08% (THI = 60), 0.44% (THI = 65), 1.7% (THI = 70), and 4.8% (THI = 75), approximately corresponding to a quadratic increase in losses. Hence, even amplifying losses for RY_F and RY_P, with animals in early lactation (<30 days) being slightly less affected. The effect of heat stress is amplified by the age of animals, whereas first lactation cows reduced milk yield by 3.0% at THI = 75, compared to increasing losses in second (4.9%), third (5.4%), and later parity (5.8%). Furthermore, animals in the first 30 (and to some extent 100) days in milk were less affected by heat stress conditions (Figure 7.B). Results of the kernel regression to estimate population-wise effects indicate systematic biases in the fitting of the lactation curves based on DIM with an estimated effect of 0.01 at around DIM 30, which should however be corrected by the inclusion of this effect (Supplementary Figure S18). Similarly, slightly lower milk yield was observed in spring and fall compared to summer and winter.

**Figure 7:**
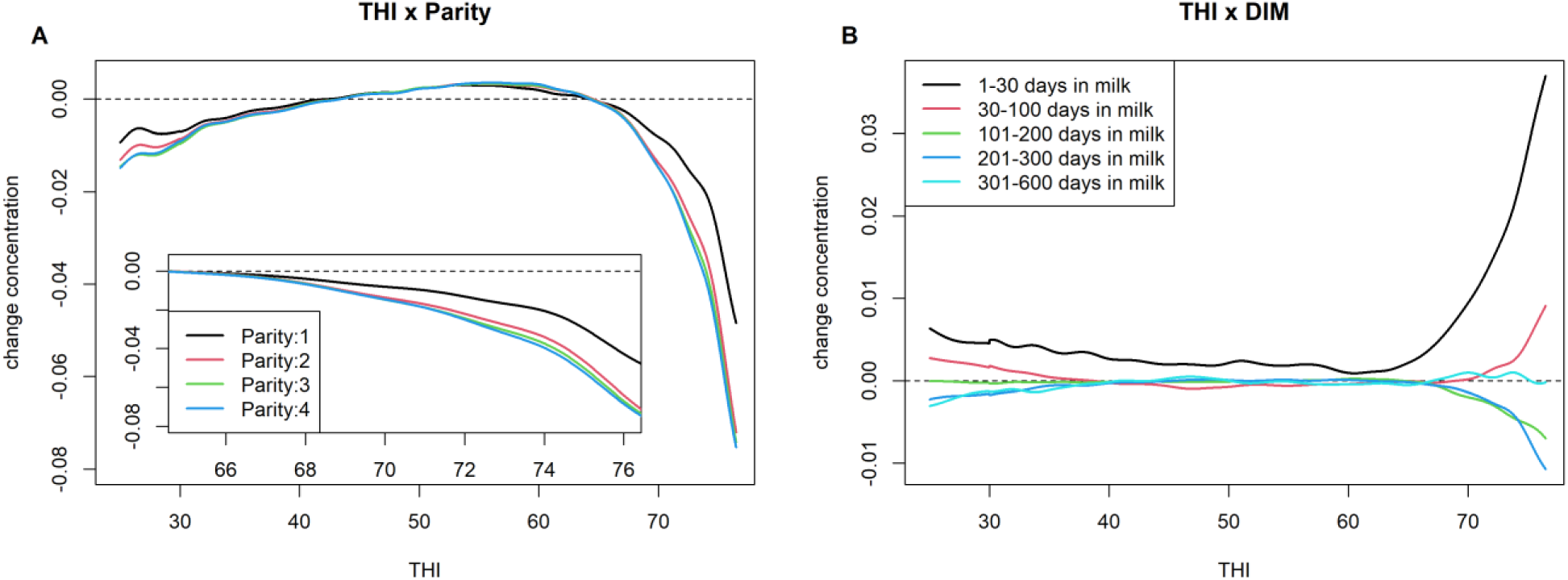
Estimated effects of THI x parity (a) and THI x DIM (b) on milk yield in percent relative to expectation based on the lactation curve.

Obtained models when scaling solid components by the animal’s own performance relative to the own lactation curve resulted in very similar models to the concentration-based models up to THI = 60 and subsequent stronger losses in both RY_F and RY_P in line with the cumulative effects seen in F% or P% and milk yield traits combined (Supplementary Figures S19 and S20).

### Individual phenotypes

Based on our newly proposed trait definition for heat tolerance traits, phenotypes for 273,135 out of 346,248 cows were derived (of which 56,505 are genotyped). When setting all records below THI = 50 to 50, the number of phenotyped cows was reduced to 271,634 as some cows had no observation associated with a THI higher than 50. In contrast, requiring a higher number of records heavily reduced the share of phenotyped animals, e.g., requiring 10 or 15 records would result in just 213,227 or 160,140 cows with phenotypes, respectively. The requirement of a higher number of records particularly affects young animals.

Estimated heritabilities for the heat tolerance trait for F% (H_F%) and P% (H_P%) were 0.046 and 0.116 (Table 2). The genetic standard deviation (gSD) for the two traits was 0.004 and 0.003, respectively (Table 2). This implies, that a cow that is one gSD superior in H_F% is expected to have 0.086% higher F% at THI = 70 than a standard cow with the same performance at THI = 50. The heritability for heat tolerance for milk yield (H_MY) was 0.095 with a gSD of 0.0010 (corresponding to an expected 0.2% higher milk yield for a one gSD superior cow at THI = 70).

**Table 2:**
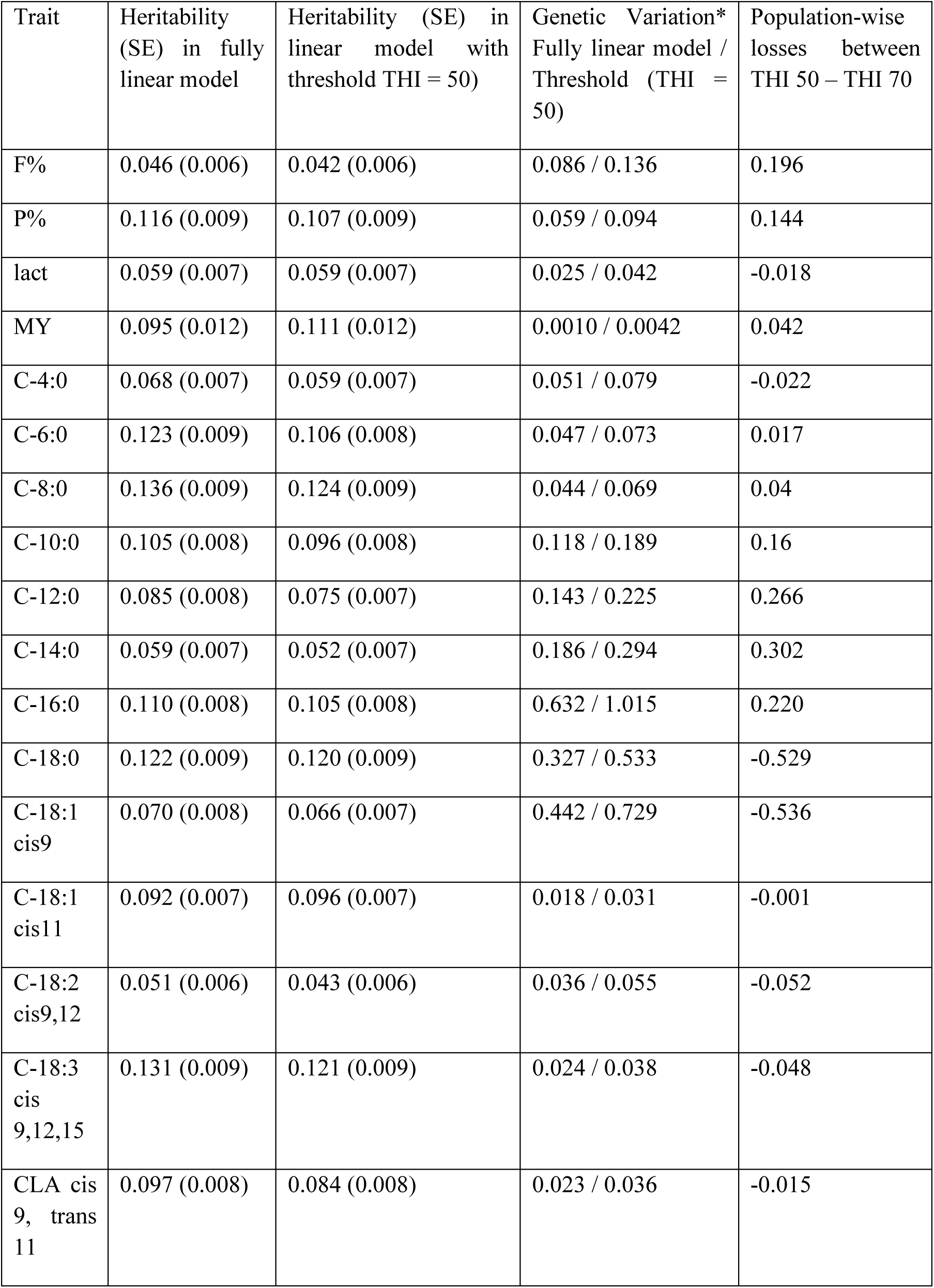
Estimated variance components for different heat tolerance traits and their genetic variation compared to population-wise effects. *The genetic variation is expressed as the genetic standard deviation * 20 to express the differences between a ”standard” cow and 1 gSD superior cow in THI = 50 and THI = 70 conditions

| Trait | Heritability (SE) in fully linear model | Heritability (SE) in linear model with threshold THI = 50) | Genetic Variation* Fully linear model / Threshold (THI = 50) | Population-wise losses between THI 50 – THI 70 |
| --- | --- | --- | --- | --- |
| F% | 0.046 (0.006) | 0.042 (0.006) | 0.086 / 0.136 | 0.196 |
| P% | 0.116 (0.009) | 0.107 (0.009) | 0.059 / 0.094 | 0.144 |
| lact | 0.059 (0.007) | 0.059 (0.007) | 0.025 / 0.042 | -0.018 |
| MY | 0.095 (0.012) | 0.111 (0.012) | 0.0010 / 0.0042 | 0.042 |
| C-4:0 | 0.068 (0.007) | 0.059 (0.007) | 0.051 / 0.079 | -0.022 |
| C-6:0 | 0.123 (0.009) | 0.106 (0.008) | 0.047 / 0.073 | 0.017 |
| C-8:0 | 0.136 (0.009) | 0.124 (0.009) | 0.044 / 0.069 | 0.04 |
| C-10:0 | 0.105 (0.008) | 0.096 (0.008) | 0.118 / 0.189 | 0.16 |
| C-12:0 | 0.085 (0.008) | 0.075 (0.007) | 0.143 / 0.225 | 0.266 |
| C-14:0 | 0.059 (0.007) | 0.052 (0.007) | 0.186 / 0.294 | 0.302 |
| C-16:0 | 0.110 (0.008) | 0.105 (0.008) | 0.632 / 1.015 | 0.220 |
| C-18:0 | 0.122 (0.009) | 0.120 (0.009) | 0.327 / 0.533 | -0.529 |
| C-18:1 cis9 | 0.070 (0.008) | 0.066 (0.007) | 0.442 / 0.729 | -0.536 |
| C-18:1 cis11 | 0.092 (0.007) | 0.096 (0.007) | 0.018 / 0.031 | -0.001 |
| C-18:2 cis9,12 | 0.051 (0.006) | 0.043 (0.006) | 0.036 / 0.055 | -0.052 |
| C-18:3 cis 9,12,15 | 0.131 (0.009) | 0.121 (0.009) | 0.024 / 0.038 | -0.048 |
| CLA cis 9, trans 11 | 0.097 (0.008) | 0.084 (0.008) | 0.023 / 0.036 | -0.015 |

Setting all THI values at colder conditions to a predefined threshold, mimicking a broken stick model, resulted in higher genetic variance and heritabilities comparable to those from the baseline models, e.g. with a threshold of THI = 50 leading to heritabilities of 0.044 (H_F%), 0.107 (H_P%), 0.111 (H_MY) and genetic standard deviations of 0.0068, 0.0047, and 0.0021 respectively (Table 2). Compared to the population-wise effects of heat stress, the relative reduction of the three heat tolerance traits based on F%, P%, and MY from THI = 50 to THI = 70 conditions would be reduced by 69%, 65%, and 10% respectively for a 1 gSD superior animal (Table 2).

Requiring a higher number of records before assigning a phenotype to the animal did increase the heritability with e.g., a minimum of 15 records (instead of 5) resulting in a heritability of 0.070 (instead of 0.046) for H_F%, however, this was only caused by a reduction of residual variance as genetic variance stayed the same or in case of H_MY even slightly reduced (Supplementary Table S1).

Estimated heritabilities for the intercepts were generally high for all milk composition traits considered with values of 0.73 and 0.74 for H_F% and H_P%, respectively. Note that these values are higher than values reported in the literature (Druet et al., 2005; Soyeurt et al., 2007), however, consider that phenotypes are not derived from a single milk sampling, and residual variances are reduced by correction for environmental effects. Thereby, what is here reported as a heritability from a comparability perspective is more in line with repeated sampling / repeatability. From this comparison perspective, estimates are in line with estimates reported for repeatability for F% and P% (National Research Council, 1988; Costa et al., 2019). The heritability for the intercept of H_MY was practically zero, as milk yield is reported relative to the individual lactation curve, resulting in the same target intercept for each animal, independent of its genetics.

As variance component estimation indicated the highest promise of trait definition that set records with THI below 50 to that threshold, all subsequent analyses on genetic correlation and GWAS primarily focus on this trait definition. Note that correlations to phenotypes calculated based on the linear model without alternating THI values were on average 0.91, therefore showing very similar correlations and GWAS peaks.

The correlations between heat tolerance traits (Figure 8.A) were similar to correlations between the different milk components themselves (Figure 8.B), e.g., with F% and P% having a residual correlation of 0.33 and a genetic correlation of 0.82, being approximately in line with estimates from Soyeurt et al. (Soyeurt et al., 2007). Furthermore, high correlations between short-chain fatty acids (C-6 to C-14) and long-chain fatty acids (C18:1 cis9 – CLA cis 9,trans11) among each other were observed, whereas correlations between the two groups were mostly negative. Correlations between H_MY and milk component-based traits were all around zero. Correlations between intercept and slope of the considered heat tolerance traits were mostly negative, with a correlation of -0.48 and -0.36 for F% and P%, indicating that animals with lower overall production levels in absolute terms have a smaller drop in performance in heat conditions.

**Figure 8:**
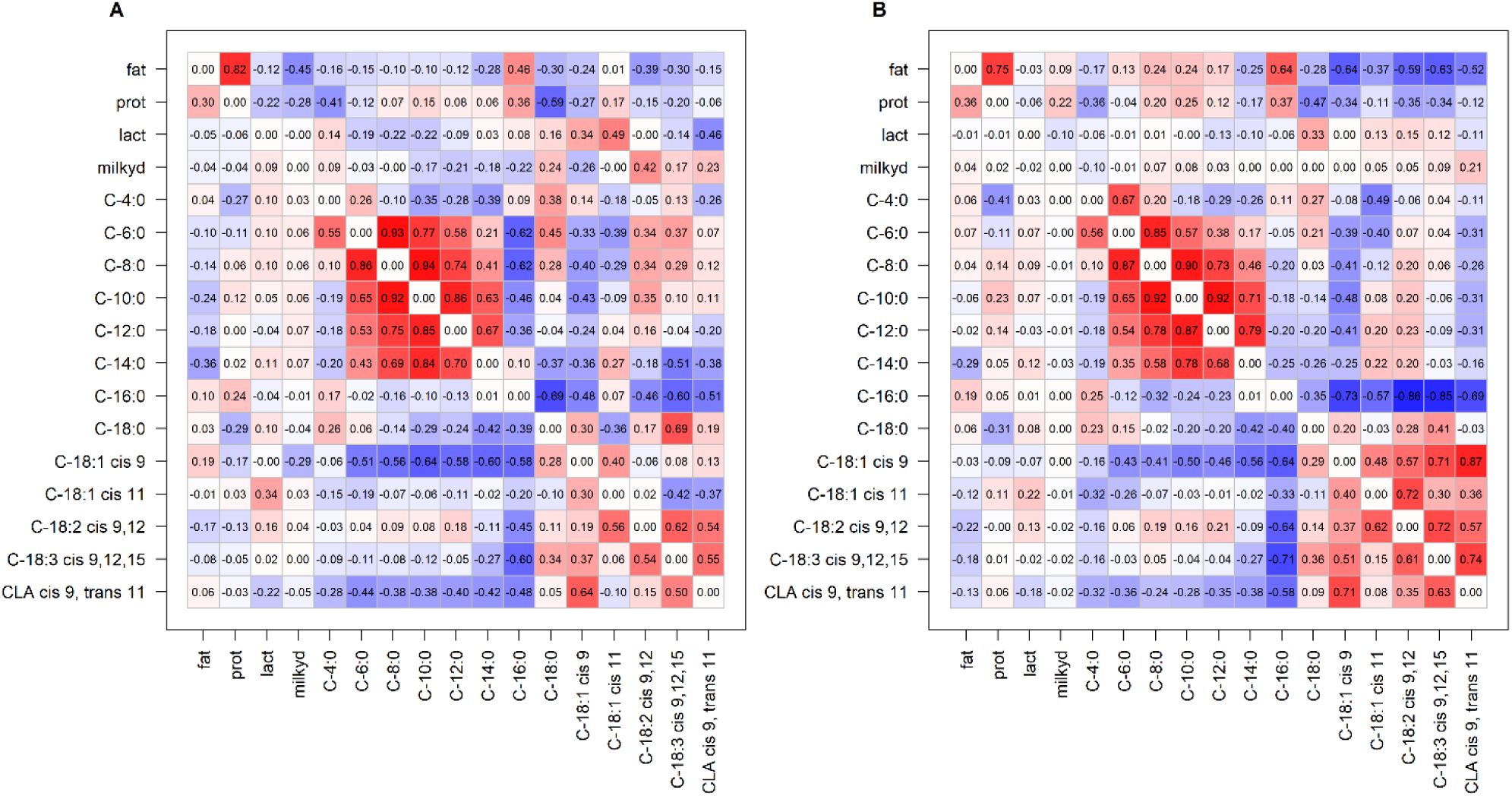
Correlations between heat tolerance traits based on milk yield and fat, protein and fatty acid concentration based for the slope (A; heat tolerance) and the intercept (B, performance at thermoneutral conditions). For milk yield in both figures the slope is used as by design the intercept has a target value of 0 for all individuals. Genetic correlations are given above the diagonal and residual correlations below the diagonal.

Correlations to commercial traits for H_MY were mostly beneficial with an overall correlation of 0.29 to the total merit index (NVI, Table 3), with particularly favorable correlations to longevity (0.33). Conversely, correlations of H_F% and H_P% to the NVI were -0.14 and -0.09 with negative correlation to resilience (-0.22 / -0.19), claw health (-0.16 / -0.12), and subclinical ketosis (-0.12 / -0.19).

**Table 3:** Estimated genetic correlations between heat tolerance traits (slope) for selected commercial traits to heat tolerance based on MY, F%, and P%. A full list of traits and correlations to individual fatty acid based heat tolerance traits is given in Supplementary Figure S21.

| Trait | H_F% | H_P% | H_MY |
| --- | --- | --- | --- |
| MY | +0.17 | +0.19 | +0.03 |
| Fat yield | -0.24 | -0.17 | +0.04 |
| Protein yield | -0.03 | -0.04 | +0.11 |
| Longevity | -0.05 | -0.01 | +0.33 |
| Fertility | -0.07 | -0.07 | +0.17 |
| Udder health | -0.01 | -0.01 | +0.12 |
| Subclinical mastitis | -0.04 | -0.02 | +0.11 |
| Subclinical ketosis | -0.12 | -0.19 | +0.15 |
| Resilience | -0.22 | -0.19 | +0.18 |
| Stability | -0.17 | -0.21 | +0.27 |
| Total Merit Index | -0.14 | -0.09 | +0.29 |

### Genome-wide association study

The use of HaploBlocker (Pook et al., 2019) resulted in the identification of 154,318 haplotype blocks, effectively tripling the number of considered variants compared to when only using a panel of SNPs.

Results for GWAS studies showed significant effects in various regions of the genome with the highest peak for most traits identified on chromosome 14 with peaks closely matching the location of DGAT1 (Prakapenka et al., 2024) (Figure 9). Note that all GWAS were also performed including DGAT1 as a fixed effect, however, with the exception of the DGAT1 region having no significant associations anymore, results stayed basically the same. Across all traits, a series of SNPs showed highly significant hits (p < 10^-15) for multiple of the 17 considered traits with the most frequent hits on BTA14 (0.1 MB, 15 hits), BTA5 (11 hits, 102.0 Mb), BTA27 (10 hits, 38.9 Mb), and BTA 19 (9 hits, 52.1Mb). A full list of those most frequently significant variant per chromosome is given in Table 4. Furthermore, special attention was given to the peak on BTA20 as it showed a significant peak for H_F%, H_P%, and H_MY (Figure 10). Of the variants with frequently observed hits, regions on BTA5 (Sigdel et al., 2019), BTA14 (Prakapenka et al., 2024), BTA19 (Bouwman et al., 2014), and BTA20 (Huson et al., 2014) have all been previously associated with milk composition or heat tolerance. Note, that the SLICK variant (Huson et al., 2014), the primary genetic variant reported on BTA20 associated with heat tolerance, is not present in the Dutch Holstein population. The SNP with the highest peak on chromosome 20 for H_F% is at 36.4 MB with a frequency of 10.4% in the population. Contrarily, the highest peak haplotype block has a frequency of 7.0% and spans 10 SNPs from 33.2 MB to 33.6 MB. Note that all these SNPs have a frequency of at minimum 27.7% in the population and only the combined sequence of allelic variants is present at 7.0% frequency.

**Figure 9:**
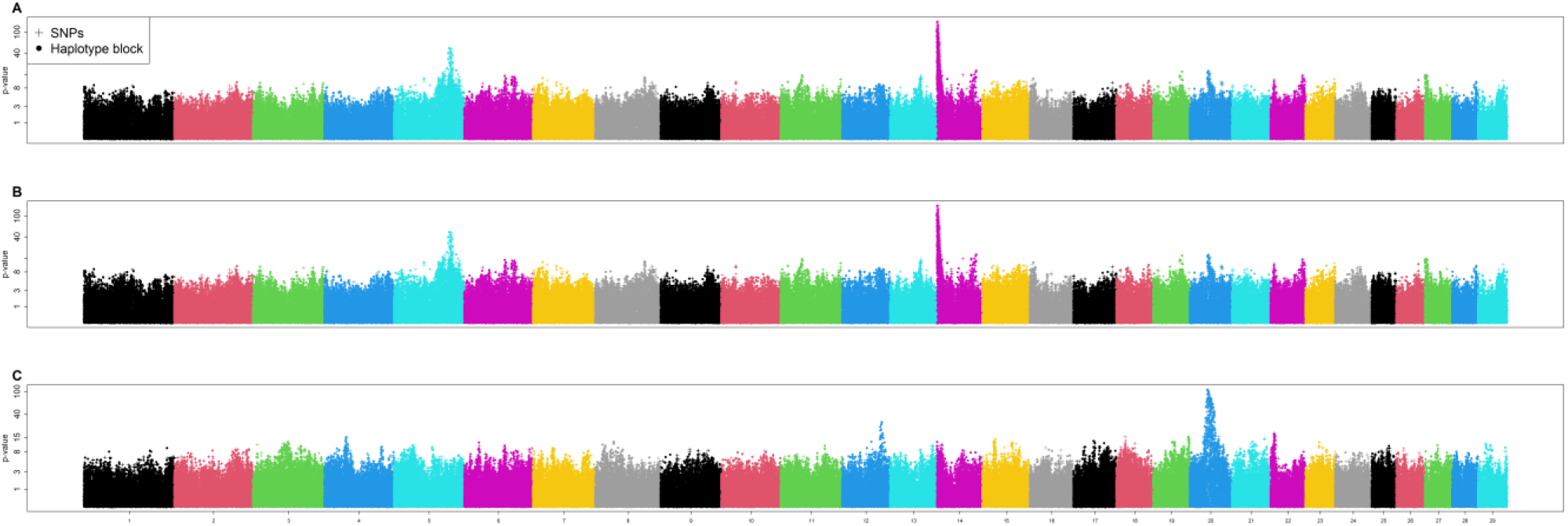
Manhattan plot based on a Genome-wide association study for the heat tolerance trait for H_F% (A), H_P% (B), and H_MY (C). The y-axes are log-scaled.

**Figure 10:**
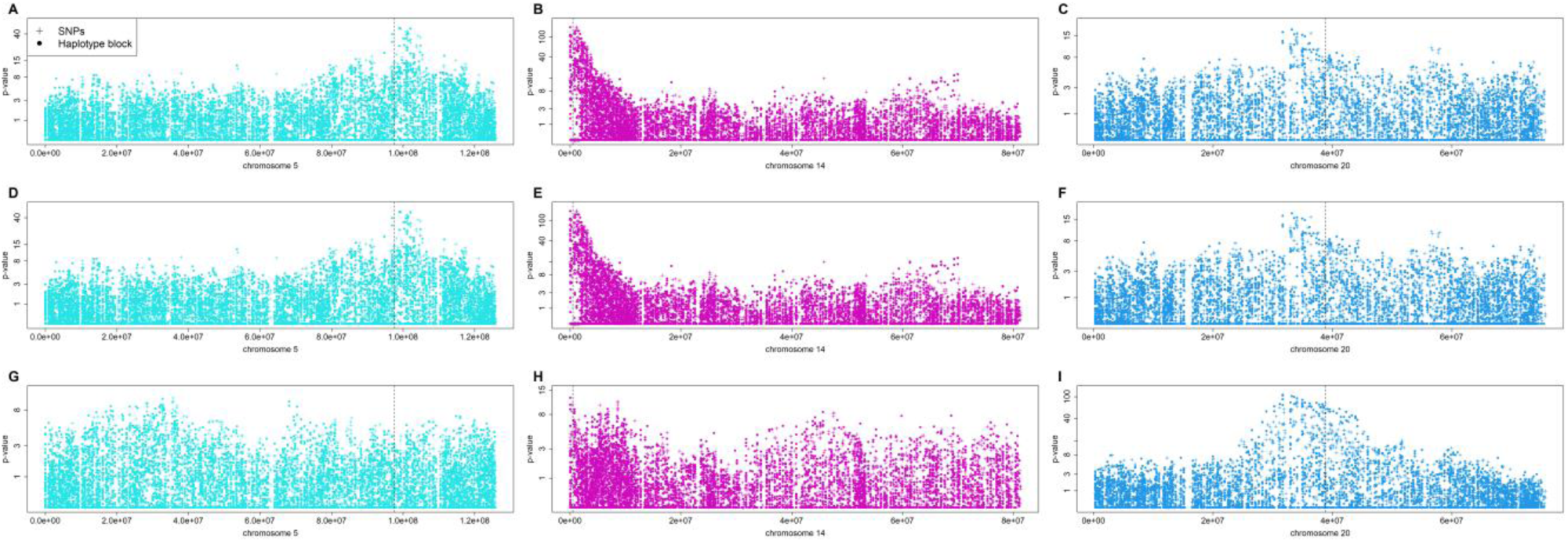
Manhattan plot based on a Genome-wide association study for the heat tolerance trait for H_F% (A,B,C), H_P% (D,E,F), and H_MY (G,H,I) on BTA5 (A,D,G), BTA14 (B,E,H), BTA20 (C,F,I). Location of CDKN1B and DUSP16 (Sigdel et al., 2019), DGAT1 (Prakapenka et al., 2024), and SLICK (Huson et al., 2014) are marked with vertical lines. The y-axes are log-scaled.

**Table 4:** Overview of SNPs / haplotype blocks with the highest number of QTLs hits (p < 10^-15) for the 17 heat tolerance traits considered.

| SNP ID | Chromosome | Location (in Mb) | Number of associations ( $p < 10^{-15}$) |
| --- | --- | --- | --- |
| Haplotype Block | 14 | 0.01 - 0.84 | 15 |
| Haplotype Block | 5 | 102.05 - 102.57 | 11 |
| ARS-BFGL-NGS-57448 | 27 | 38.88 | 10 |
| BTA-45758 | 19 | 52.1 | 9 |
| SNP_X14710_1740 | 11 | 107.17 | 7 |
| Haplotype Block | 26 | 9.35 - 9.75 | 7 |
| ARS-BFGL-NGS-39003 | 15 | 65.04 | 7 |
| SNP_BES11_Contig346_1209 | 13 | 64.81 | 6 |
| Haplotype Block | 2 | 130.49 - 131.78 | 6 |
| MARC_33120_400 | 17 | 31.35 | 5 |
| Haplotype Block | 3 | 16.22 - 16.85 | 5 |
| Haplotype Block | 10 | 6.13 - 6.35 | 4 |
| Haplotype Block | 20 | 31.69 - 34.04 | 3 |
| Haplotype Block | 9 | 105.68 - 108.07 | 3 |
| ARS-BFGL-NGS-100347 | 25 | 28.56 | 3 |
| BTB-01958090 | 16 | 64.21 | 3 |
| rs29024685 | 6 | 88.54 | 3 |
| Hapmap40784-BTA-66134 | 29 | 10.41 | 3 |
| AAFC03097715_86987 | 1 | 145.85 | 3 |
| BTB-01381318 | 23 | 52.27 | 2 |
| Haplotype Block | 18 | 57.99 - 60.29 | 2 |
| Haplotype Block | 12 | 69.42 - 70.85 | 2 |
| Haplotype Block | 7 | 47.1 - 58.16 | 1 |
| BTA-64106 | 28 | 34.11 | 1 |
| ARS-BFGL-NGS-38990 | 4 | 101.67 | 1 |
| Haplotype Block | 22 | 61.06 - 61.33 | 1 |
| Haplotype Block | 21 | 67.55 - 68.17 | 1 |
| Haplotype Block | 24 | 53.88 - 54.18 | 1 |
| --- | 8 | --- | 0 |

On a phenotypic level, carriers of the peak SNP have a higher heat tolerance in terms of H_F% (allele count 0: -0.0011; allele count 1: +0.0033; allele count 2: +0.0079; Figure 11). Similarly, animals carrying the peak haplotype also have higher heat tolerance (haplotype count 0: -0.001; haplotype count 1: +0.0053; haplotype count 2: +0.0085). However, animals that carry the peak SNP and not the haplotype show a substantially lower change in heat tolerance (allele count 0: -0.0011; allele count 1: +0.0001; allele count 2: +0.0025 with haplotype count 0). As these carriers are part of various other haplotype blocks (Figure 12), it seems likely that the SNP was only identified due to its high LD with the peak haplotype block but is less linked to the causal variant than the haplotype block. When comparing haplotype blocks in chromosome 20 along the estimated effect for H_F% (Figure 13), the highest effects are estimated for blocks in strong linkage with the peak haplotype with only minor differences between alternative haplotype blocks in the genome segment. Overall, haplotype blocks improved resolution noticeably with blocks frequently having the lowest p-value in the peak regions including regions on BTA5, BTA14, and BTA20 (Figure 14 and Table 4) and provide a visualization tool to assist with fine-mapping.

**Figure 11:**
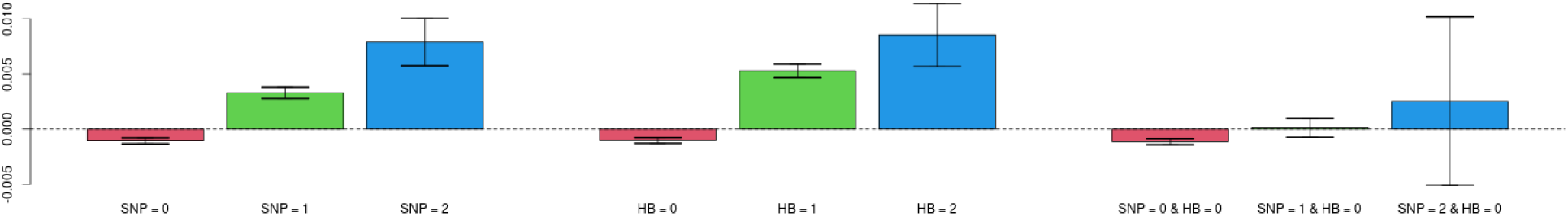
Average phenotypic level for H_F% depending on allelic variant on peak SNP and haplotype count at peak haplotype block on chromosome 20. The confidence band (95%) is calculated for the estimated mean phenotype.

**Figure 12:**
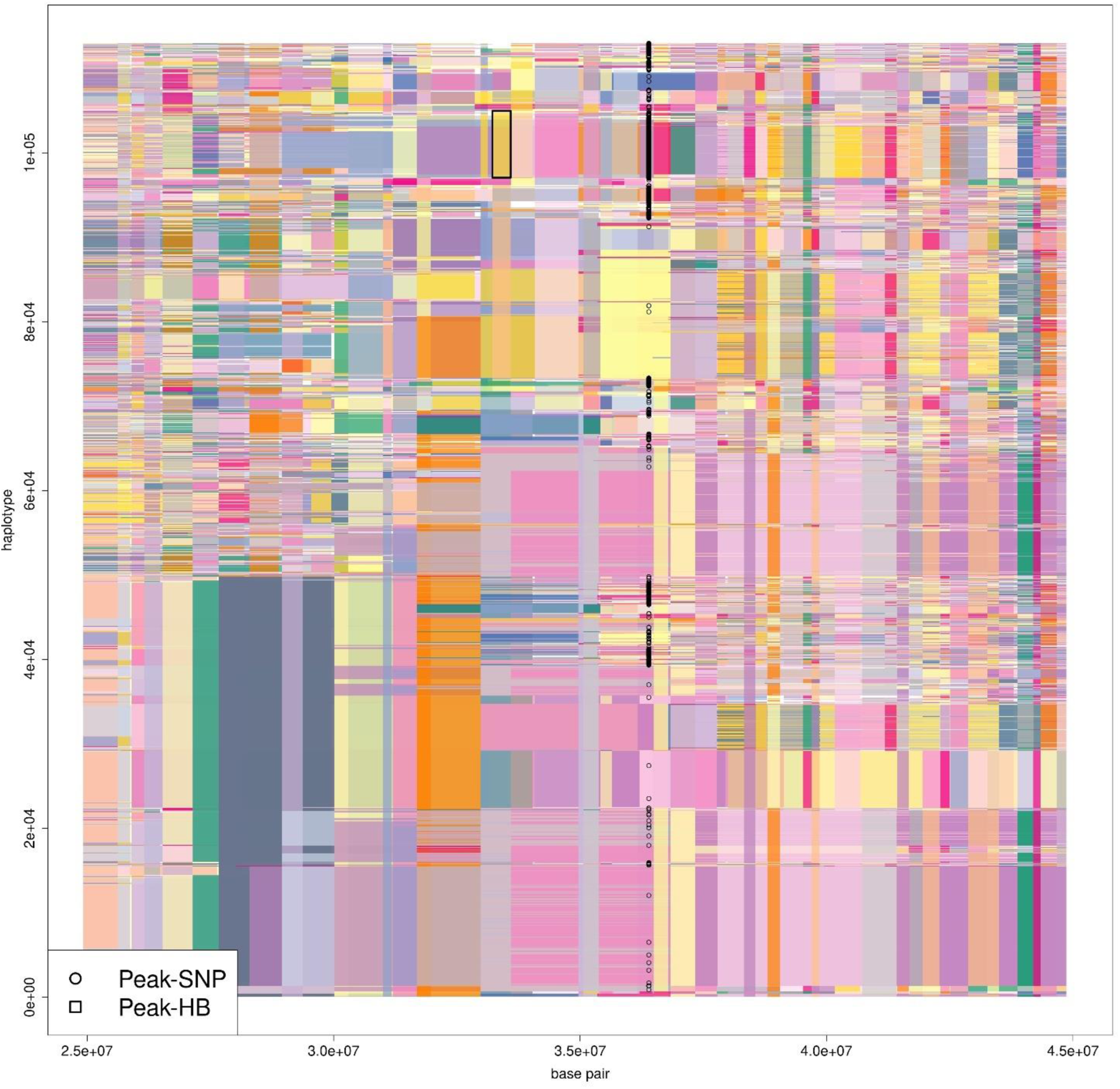
Haplotype based visualization of individuals with a phenotype for H_F% and available genomic data. Haplotypes are sorted according to their similarity at 33.3 Mb according to Pook et al. (2019). Haplotypes with the beneficial variant at the peak SNP and haplotype block are indicated in black.

**Figure 13:**
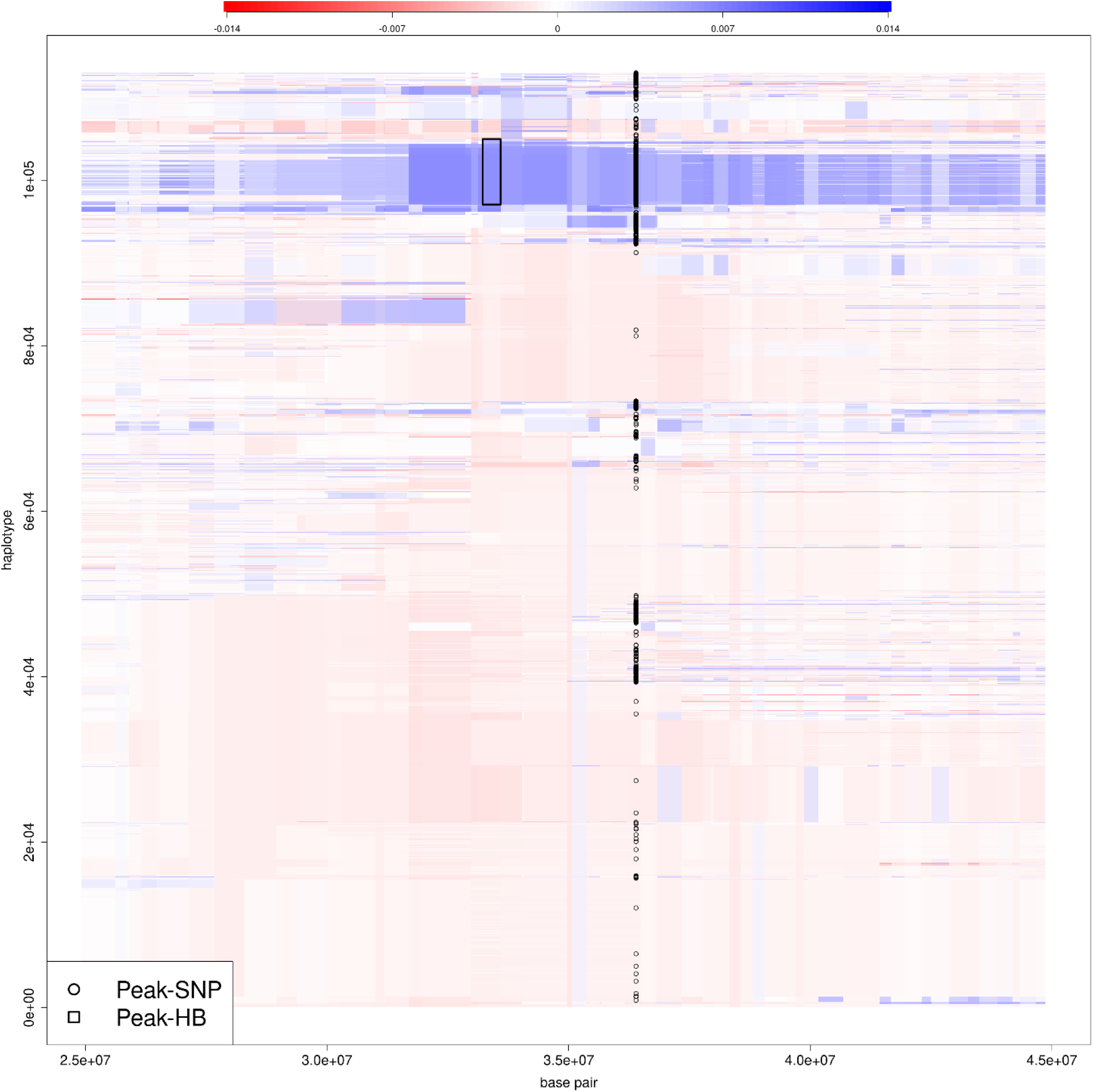
Haplotype based visualization of individuals with a phenotype for H_F% and available genomic data. Haplotypes are sorted according to their similarity at 33.3 Mb according to Pook et al. (2019). The color of blocks is chosen according to the estimated effect of each haplotype block based on phenotypic data with high heat tolerance indicated in blue and low heat tolerance indicated in red. Haplotypes with the beneficial variant at the peak SNP and haplotype block are indicated in black.

**Figure 14:**
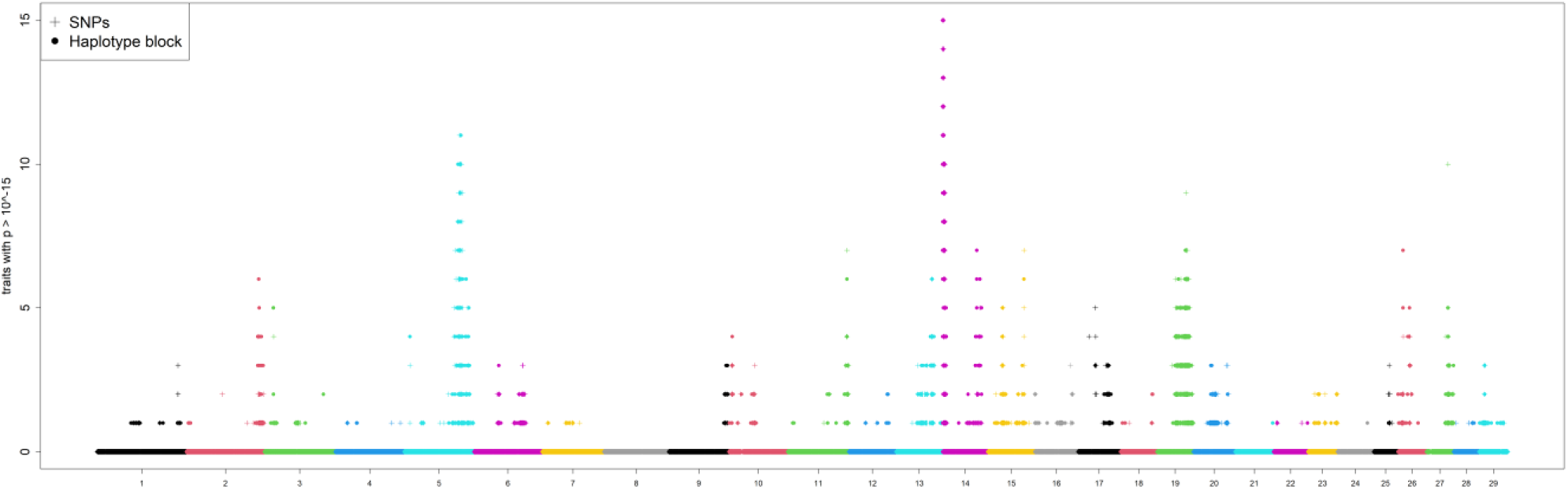
Number of heat tolerance traits each SNP / haplotype has a p-value < 10^-15 for the 17 considered milk composition traits (using the slope estimated when setting records to THI < 50 to 50).

### Computing time

On the computational side, the entire pipeline for population-wise effects took 41 minutes with a peak memory usage of 35 GB using a single core of an Intel Xeon Gold, 2.1 GHz processor per trait. Both memory and computing times scale approximately linearly in the number of records included. The subsequent derivation of phenotypes required neglectable computing resources, completing in 3.3 minutes with 1.6 GB peak memory for processing all traits combined. Note here that our pipeline mostly uses basic R-functions like *ksmooth()* and *lm()* (R Core Team, 2017) and is by no means optimized for the computational efficiency of the concrete application at hand.

## Discussion

In this work, we are providing a comprehensive new pipeline for the estimation of both individual-based and population-wise effects of heat based on repeated measurements as an alternative to reaction-norm models. Our proposed approach allows more sophisticated models to include non-linear effects or interactions between parameters, such as THIxDIM interactions. While applied on milk yield and milk composition traits in this study, the proposed population-wise models can be applied in the same way to other traits or species without restrictions, whereas the suggested novel traits require the availability of longitudinal data.

### Population-wise models

Our results highlight the impact of weather conditions on dairy cattle indicated by a linear reduction of F% and P% across the entire spectrum of considered THI values. In contrast, milk yield itself was relatively stable till a THI of 60, however, with rising THI values, production losses increased quadratically. Animals in higher parity were more strongly affected by heat, as also reported by Aguilar et al. (2010). In contrast to Aguilar et al. (2010), our results suggest a noticeable impact of heat for first parity cows. As only data from Dutch farms with a temperate maritime climate were considered in this study, effects could only be reliably estimated up till THI = 75, where milk yield was already reduced by 5%. Extending the observed quadratically increasing losses suggests a reduction of milk yield by 15-20% at THI = 80. In view of climate change (Cheng et al., 2022) such levels of heat stress must be expected in the Netherlands in the future and are common in other parts of the world today. Practically, we would expect losses to be even higher as F% and P% are also expected to reduce. Also, in our study, the most critical cases of heat stress in which an animal did not produce any milk were removed from the analysis, as they were very rare in our specific data and we had no health data to correct for other potential reasons of complete milk production losses. Lastly, we expect farmers to adapt feeding and husbandry conditions in extreme weather conditions, leading to a potential underestimation of the effects of heat stress (Johnson, 1987; D’Emilio et al., 2017; Ríus, 2019).

Note that the interpretation of the individual effects of any population-wise models should be done with caution as individual parameters can be highly interdependent, leading to multicollinearity (Alin, 2010). For example, the season effect in summer months showed the lowest expected F% and P%, potentially leading to a further underestimation of the effect of heat stress. We also considered models that include other, more detailed weather data, such as the number of hours in a day above a certain THI threshold, maximum THI / temperature, and air pressure (Misztal et al., 2024). Whereas goodness-of-fits was basically not affected for thermoneutral conditions (35 < THI < 65), the absolute size of residual effects in extreme weather situations was reduced, thereby indicating a better fit and/or overfitting. For the reason of interpretability and to reduce overfitting, the finally used models in this study were kept relatively simple.

Although THI is not a perfect metric for determining the overall heat load, one of its core advantages is its simplicity, easily allowing for an interpretation of the estimated effects and subsequent fitting of a regression line. For more complex models, we propose to aggregate all effects associated with climatic conditions to derive an overall weather effect for each farm-by-day combination that should provide a more advanced rating of the severity of heat stress on a given day. However, since seasonal effects will also include other effects like the change in diet over the year, isolating the effect of heat stress will not be fully possible.

### Breeding for heat tolerance

In regard to the definition of our newly suggested heat tolerance traits, our suggested approach has similarities to the slope in a reaction-norm model (Su et al., 2006; Aguilar et al., 2010). However, in terms of its use in breeding, we are proposing the use of the estimated slope as the trait to breed for, acting as a residual trait. Contrary to this, although the slope is an output from a reaction-norm model the realized estimated breeding values at different THI values are usually proposed for usage, as heritability for the slope in reaction-norm models is often not significantly different from zero (Copley, 2024), leading to high correlations to between traits at different THI levels while our residual trait if at all showed negative correlation to overall production levels (intercepts).

As repeated records of a cow are combined into a single phenotypic record, our proposed approach will result in reduced computational load compared to reaction-norm models, when applied in routine genetic evaluations. As shown above, the computation of pre-corrected phenotypes based on a population-wise models is relatively straightforward, even for large datasets,. Furthermore, the overall flexibility to include additional effects is greatly increased.

It should still be noted that this comes with the limitation that longitudinal data for an animal is required in our approach. Note that from a goodness-of-fit perspective, an iterative fitting procedure is inferior compared to a joint / all-in-one model (Holland and Piepho, 2024), at least if both models are able to incorporate the same parameters and their effect structures. On the other hand, a downstream application like GWAS in our approach can be performed directly on phenotypes instead of estimated breeding values or deregressed proofs, thereby being conceptionally better suited than application on a value derived based on SNP effects (Ekine et al., 2014; Holland and Piepho, 2024).

A limitation of our approach is that the reliability of the phenotype of an animal is currently not accounted for in the prediction model. As such, cows with a higher number of observations will have more reliable phenotypes, paired with phenotypes not being derived based on a single record but longitudinal data. This in turn also leads to higher heritability estimates when only including animals with a high number of individual records – as genetic variance did not increase and fewer phenotypes for young animals will be available, we do however not expect higher prediction accuracies for young animals.

From a methodological perspective, this does not pose a conceptual issue but is an important consideration when interpreting the estimated heritability. We explored integrating the 𝑅^2^ values from the linear regression in our models as an indicator of the reliability of the individual phenotypes, however, values overall were low with 𝑅^2^ < 0.01 for 95% of all samples for H_MY (Supplementary Figure S19) as the variance of production levels are much higher than the isolated effect of weather conditions (Figure 3).

A potential solution for differences in the reliability of individual phenotypes could be to process each lactation of an animal as a separate phenotypic record. This should provide similar reliabilities between individual phenotypes, while ensuring that each phenotype is derived based individual records from most seasons of the year. Thereby, also avoiding potential biases introduces by partial lactation or frequent re-ranking animals.

The low values for 𝑅^2^again highlight the need to use longitudinal data to improve the accuracy of prediction models, going even beyond the nowadays commonly used monthly test day recording scheme and instead considering weekly or even more ideally daily or per-milking recording. Although scaling of reaction-norm models is linear in the number of records for many implementations solving mixed model equations, computing times with billions of records are expected to be critical while our pipeline is able to summarize repeated records of the same animal efficiently, thereby resulting in subsequent prediction models that only include a single record (or one per lactation).

In terms of the overall potential for breeding, all our heat tolerance traits showed noticeable variance. However, for practical breeding, it is probably required to combine individual traits into a joint trait (complex) to avoid a rapid increase in the number of traits considered. In this regard, H_F% and H_P% show a high correlation, making a combination of these two easily possible. As genetic variances in both H_F% and H_P% are high relative to the population-wise effects of heat (69% / 65%), this indicated that with relatively little breeding effort reduced concentration due to heat can be compensated. Milk yield remained stable up to a THI of 60. However, at higher THI levels, production losses increased substantially. Despite substantial genetic variation in H_MY, a 1 gSD improvement would only offset 11% of the losses. Nonetheless, as there were no negative correlations to milk, protein, or fat yield, paired with favorable correlations to other production traits (0.29 correlation to NVI) the use for breeding should come at limited risk.

As there is currently no economic benefit in changing individual fatty acid concentration, there is no direct benefit of the use of the heat tolerance traits for individual fatty acids, however, these traits could still be beneficial for breeding purposes as indicator traits for heat stress with H_C-16:0 and H_C-18:0 showing significant correlation to resilience and H_C-18:3cis9,12,15 to longevity (Figure 9). Furthermore, association with other heat-related traits like rectal temperature, respiratory rate, or activity (Li et al., 2020; Yan et al., 2021) should be assessed to evaluate the potential to replace or complement costly recording schemes for such traits by the use of already available MIR data.

### Between heat resilience and tolerance

On a conceptional level, further consideration needs to be that the here-defined heat tolerance traits primarily focus on which animals are able to maintain their production levels the best under heat conditions. However, reduction of production levels can sometimes be the only way for the animal to cope with the extreme stress situation to avoid more severe reactions with the most extreme reaction being the likelihood of death (West, 2003), commonly referred to as the heat resilience of an animal (Misztal et al., 2024). In practical breeding, yield-based traits as proposed in this work should be complemented by health records to capture both heat tolerance - ensuring stable production - and heat resilience - reducing the risk of morbidity and mortality - within the breeding objective (Poppe et al., 2022; Misztal et al., 2024). Our results here indicate that cows that reduce their production levels by decreasing F% and P% tend to have worse resilience, while a smaller reduction of milk yield in heat conditions showed favorable correlations to health and resilience traits.

Improving the current population to a level of suitability in more challenging climatic conditions from genetic variation within the Dutch Holstein population will be extremely challenging. At the same time, this also highlights the need to look into alternative strategies when aiming to improve the heat tolerance of the Dutch Holstein population. Exemplary strategies could include the use of crossbreeding to introduce genetic material from a locally adapted breed or the introgression of such diversity into the breeding population (Hoffmann, 2013; Galukande et al., 2013). Although practically not possible for production animals in Europe, the downsides of introgression could be bypassed by genome-editing (Jinek et al., 2012), as already done in research with the introduction of the SLICK mutation into Holstein via the use of CRISPR-Cas9 (Cuellar et al., 2024). Before implementing such procedures, a functional understanding of the trait is however needed, particularly to ensure that a variant is actually causal (Simianer et al., 2018), as we for example identified GWAS peaks on BTA20, in close proximity to the SLICK gene (Littlejohn et al., 2014), with the SLICK variant not even present in the Dutch Holstein population. Hence, there might be further causal genes in the region that need to be accounted for. Furthermore, we observed lower overall production levels in cows carrying some of the QTLs identified in this study, introducing a further potential source of confounding by identifying lower overall production level cows as particularly heat tolerant as they just have less to lose.

### Haplotype blocks for association studies

The use of haplotype blocks (Pook et al., 2019) in a GWAS provides a powerful tool to increase the density of variants considered to add further impactful variation to a dataset, thereby increasing the power of the GWAS. As linkage and haplotype analysis (Gabriel et al., 2002; Bulik-Sullivan et al., 2015) are common analysis steps based on the results of the GWAS study, this allows to combine processing steps and achieves higher mapping precision before proceeding with further downstream analysis. Haplotype blocks have not only shown beneficial within a GWAS, but also as a visualization tool to identify phenotypic differences between subpopulations, assess local linkage, and use in fine-mapping. For the identification of causal variants, molecular analysis is required, however, quantitative / statistical analysis by haplotype blocks is a powerful tool to assist such molecular analysis by identifying candidate regions and individuals (Gusev et al., 2016; McLaren et al., 2016).

## Conclusions

Overall, we conclude that heat tolerance is an important trait in dairy cattle both from an animal welfare and production perspective and has substantial genetic variation to be utilized in breeding. Our newly developed pipeline provides a tool to computationally efficiently integrate sophisticated methodology to analyze the impact of heat stress on a population-wise level and subsequently define novel heat tolerance traits to be used for breeding based on traits with repeated measurements, such as milk production data. However, particularly in regard to heat tolerance in regard to milk yield, breeding efforts need to be complemented by other management approaches such as cooling, shading, and nutrition. To further increase genetic gain the integration of crossbreeding or intercrossing to a more adapted breed would further ways to enhance tolerance and help animals to cope with more extreme climatic conditions than what is expected in the temperate maritime climate of the Netherlands.

## Supplemental Material

Supplemental material is available at FigShare: https://figshare.com/articles/journal_contribution/Supplementary_Material_for_Pook_et_al_2025/289122 77 Doi: 10.6084/m9.figshare.28912277

Figure S1: Obtained prediction accuracies based on the PLSR model with pre-processed MIR spectra input based on the number of principle components included in the prediction model.

Figure S2: Average prediction accuracy across the 12 considered fatty acids depending on the pre-processing and the number of principle components included in the prediction model.

Figure S3-S15: Estimated effects of THI x parity (A), THI x DIM (B), season (C), and DIM (D) on lactate concentration and the different individual fatty acids in milk.

Figure S16: Estimated effects of THI (A), season (C), and DIM (D) on fat percentage in milk based on separate datasets, split by parity.

Figure S17: Estimated THI x DIM effect using all data (A), records from first (B), second (C), third (D), forth or higher (E) parity cows.

Figure S18: Estimated effects of season (A), and DIM (B) on milk yield in percent relative to expectation based on the lactation curve.

Figure S19: Estimated effects of THI x parity (A), THI x DIM (B), season (C), and DIM (D) on relative yield of fat that was calculated based on fat percentage and relative milk yield.

Figure S20: Estimated effects of THI x parity (A), THI x DIM (B), season (C), and DIM (D) on relative yield of protein that was calculated based on protein percentage and relative milk yield.

Figure S21: Estimated genetic correlations between heat tolerance traits (slope) and commercial traits.

Figure S22: 𝑅^2^ values for the linear regression models to derive heat tolerance trait for H_F% (A), H_P% (B), H_MY (C).

Table S1: Estimated variance components for different heat tolerance traits and their genetic variation compared to population-wise effects, when requiring at high number of records per cows to call a phenotype.

File S1: Overview of the estimation procedure for fatty acid concentration via partial least squares regression.

## Notes

This study was financially supported by the Horizon Europe project via the project Re-Livestock (GA No. 01059609), the Dutch Ministry of Economic Affairs (TKI Agri & Food Project LWV20054) and the Breed4Food partners CRV (Arnhem, the Netherlands), Hendrix Genetics (Boxmeer, the Netherlands), and Topigs Norsvin (Den Bosch, the Netherlands).

We acknowledge support, feedback, and discussions with Breed4Food partners in work package 3 (Phenotyping inferface) an ReLivestock partners in work package 3 (Re-Breeding Livestock).

During the preparation of this work, the authors used ChatGPT (GPT-4o) to assist with language refinement and stylistic improvements. All content was reviewed and edited by the authors, who take full responsibility for the final version of the manuscript.

MLP, IA, LZ, and CO are employed at the commercial dairy cattle breeding company CRV (www.crv4all.nl). The authors declare that this affiliation did not influence the study design, data analysis, or interpretation of results.

## Nonstandard abbreviations used

THI: temperature humidity index
GxE: genotype by environment
MIR: mid-infrared
AMS: automated milking systems
F%: fat percentage
P%: protein percentage
RY_F: relative yield of fat
RY_P: relative yield of protein
MY: milk yield
H_F%: heat tolerance for fat percentage
H_P%: heat tolerance for protein percentage
H_MY: heat tolerance for milk yield
T: temperature in Celsius
RH: relative humidity
NW: Nadaraya-Watson estimator
GWAS: genome-wide association study
MACE: multiple across-country evaluation
gSD: genetic standard deviation
NVI: total merit index
GPS: global positioning system
PLSR: partial least squares regression

## References

Aguilar, I., I. Misztal, and S. Tsuruta. 2010. Genetic trends of milk yield under heat stress for US Holsteins. Journal of Dairy Science 93(4):1754–1758.

Alin, A. 2010. Multicollinearity. Wiley interdisciplinary reviews: computational statistics 2(3):370–374.

Barrett, J. C., B. Fry, J. Maller, and M. J. Daly. 2005. Haploview: analysis and visualization of LD and haplotype maps. Bioinformatics 21(2):263–265.

Bouwman, A. C., M. H. Visker, J. M. van Arendonk, and H. Bovenhuis. 2014. Fine mapping of a quantitative trait locus for bovine milk fat composition on Bos taurus autosome 19. Journal of Dairy Science 97(2):1139–1149.

Bovine Genome Sequencing and Analysis Consortium, C. G. Elsik, R. L. Tellam, K. C. Worley, R. A. Gibbs, D. M. Muzny, G. M. Weinstock, D. L. Adelson, E. E. Eichler, and L. Elnitski. 2009. The genome sequence of taurine cattle: a window to ruminant biology and evolution. Science 324(5926):522–528.

Brockmann, M., T. Gasser, and E. Herrmann. 1993. Locally adaptive bandwidth choice for kernel regression estimators. Journal of the American Statistical Association 88(424):1302–1309.

Browning, B. L., X. Tian, Y. Zhou, and S. R. Browning. 2021. Fast two-stage phasing of large-scale sequence data. The American Journal of Human Genetics 108(10):1880–1890.

Bulik-Sullivan, B. K., P.-R. Loh, H. K. Finucane, S. Ripke, J. Yang, Schizophrenia Working Group of the Psychiatric Genomics Consortium, N. Patterson, M. J. Daly, A. L. Price, and B. M. Neale. 2015. LD Score regression distinguishes confounding from polygenicity in genome-wide association studies. Nature Genetics 47(3):291–295.

Butler, D. G., B. R. Cullis, A. R. Gilmour, and B. J. Gogel. 2009. ASReml-R reference manual. The State of Queensland, Department of Primary Industries and Fisheries, Brisbane.

Calus, M. L. P., and R. F. Veerkamp. 2003. Estimation of environmental sensitivity of genetic merit for milk production traits using a random regression model. Journal of Dairy Science 86(11):3756–3764.

Carabaño, M. J., M. Ramón, C. Díaz, A. Molina, M. D. Pérez-Guzmán, and J. M. Serradilla. 2017. Breeding and genetics symposium: Breeding for resilience to heat stress effects in dairy ruminants. A comprehensive review. Journal of animal science 95(4):1813–1826.

Cheng, M., B. McCarl, and C. Fei. 2022. Climate change and livestock production: a literature review. Atmosphere 13(1):140.

Copernicus Climate Change Service (C3S). 2024. Climate Indicators. Accessed Jan 14, 2025. https://climate.copernicus.eu/.

Copley, J. P. 2024. Genotype by environment interaction for beef cattle fertility traits in northern Australia. https://espace.library.uq.edu.au/view/UQ:0745c1c.

Copley, J. P., B. J. Hayes, E. M. Ross, S. Speight, G. Fordyce, B. J. Wood, and B. N. Engle. 2024. Investigating genotype by environment interaction for beef cattle fertility traits in commercial herds in northern Australia with multi-trait analysis. Genetics Selection Evolution 56(1):70.

Costa, A., N. Lopez-Villalobos, G. Visentin, M. de Marchi, M. Cassandro, and M. Penasa. 2019. Heritability and repeatability of milk lactose and its relationships with traditional milk traits, somatic cell score and freezing point in Holstein cows. Animal 13(5):909–916.

Cuellar, C. J., T. F. Amaral, P. Rodriguez-Villamil, F. Ongaratto, D. O. Martinez, R. Labrecque, Losano, João D de Agostini, E. Estrada-Cortés, J. R. Bostrom, and K. Martins. 2024. Consequences of gene editing of PRLR on thermotolerance, growth, and male reproduction in cattle. FASEB BioAdvances.

D’Emilio, A., S. M. C. Porto, G. Cascone, M. Bella, and M. Gulino. 2017. Mitigating heat stress of dairy cows bred in a free-stall barn by sprinkler systems coupled with forced ventilation. Journal of agricultural engineering 48(4):190–195.

Donnelly, K. P. 1983. The probability that related individuals share some section of genome identical by descent. Theoretical population biology 23(1):34–63. 10.1016/0040-5809(83)90004-7.

Druet, T., F. Jaffrézic, and V. Ducrocq. 2005. Estimation of genetic parameters for test day records of dairy traits in the first three lactations. Genetics Selection Evolution 37(4):257.

Ekine, C. C., S. J. Rowe, S. C. Bishop, and D.-J. de Koning. 2014. Why breeding values estimated using familial data should not be used for genome-wide association studies. G3: Genes, Genomes, Genetics 4(2):341– 347.

Falconer, D. S. 1952. The problem of environment and selection. The American Naturalist 86(830):293–298.

Gabriel, S. B., S. F. Schaffner, H. Nguyen, J. M. Moore, J. Roy, B. Blumenstiel, J. Higgins, M. DeFelice, A. Lochner, and M. Faggart. 2002. The structure of haplotype blocks in the human genome. Science 296(5576):2225–2229.

Galukande, E., H. Mulindwa, M. Wurzinger, R. Roschinsky, A. O. Mwai, and J. Sölkner. 2013. Cross-breeding cattle for milk production in the tropics: achievements, challenges and opportunities. Animal Genetic Resources/Resources génétiques animales/Recursos genéticos animales 52:111–125.

Garner, J. B., M. L. Douglas, O. Williams, SR, W. J. Wales, L. C. Marett, T. T. Nguyen, C. M. Reich, and B. J. Hayes. 2016. Genomic selection improves heat tolerance in dairy cattle. Scientific Reports 6(1):34114.

Gote, M. J., I. Adriaens, M. Ceccarelli, L. D. Anvers, D. Meuwissen, and B. Aernouts. 2024. CowBase-A Library for Dairy Farm Data Handling and Curation in Python. 75th Annual Meeting of the European Federation of Animal Science, Florence, Italy.

Gusev, A., A. Ko, H. Shi, G. Bhatia, W. Chung, B. W. Penninx, R. Jansen, E. J. C. de Geus, D. I. Boomsma, and F. A. Wright. 2016. Integrative approaches for large-scale transcriptome-wide association studies. Nature Genetics 48(3):245–252.

Hammami, H., J. Vandenplas, M.-L. Vanrobays, B. Rekik, C. Bastin, and N. Gengler. 2015. Genetic analysis of heat stress effects on yield traits, udder health, and fatty acids of Walloon Holstein cows. Journal of Dairy Science 98(7):4956–4968.

Hassanpour, A., J. Geibel, H. Simianer, and T. Pook. 2023. Optimization of breeding program design through stochastic simulation with kernel regression. G3: Genes, Genomes, Genetics:jkad217.

Henderson, C. R. 1975. Best linear unbiased estimation and prediction under a selection model. Biometrics:423–447.

Herbut, P., G. Hoffmann, S. Angrecka, D. Godyń, F. M. C. Vieira, K. Adamczyk, and R. Kupczyński. 2021. The effects of heat stress on the behaviour of dairy cows–a review. Annals of Animal Science 21(2):385–402.

Hoffmann, I. 2013. Adaptation to climate change–exploring the potential of locally adapted breeds. Animal 7(s2):346–362.

Holland, J. B., and H.-P. Piepho. 2024. Don’t BLUP Twice. G3: Genes, Genomes, Genetics 14(12):jkae250.

Huson, H. J., E.-S. Kim, R. W. Godfrey, T. A. Olson, M. C. McClure, C. C. Chase, R. Rizzi, A. M. P. O’Brien, C. P. van Tassell, and J. F. Garcia. 2014. Genome-wide association study and ancestral origins of the slick-hair coat in tropically adapted cattle. Frontiers in genetics 5:101.

ICAR. 2023. Section 2 - Guidelines for Dairy Cattle Milk Recording. Accessed Mar 29, 2025. https://www.icar.org/Guidelines/02-Overview-Cattle-Milk-Recording.pdf?

Jinek, M., K. Chylinski, I. Fonfara, M. Hauer, J. A. Doudna, and E. Charpentier. 2012. A programmable dual-RNA–guided DNA endonuclease in adaptive bacterial immunity. Science 337(6096):816–821.

Johnson, P. E. 1987. Practical Heat-Stress Management.

Jordan, E. R. 2003. Effects of heat stress on reproduction. Journal of Dairy Science 86:E104–E114.

Kipp, C., K. Brügemann, T. Yin, K. Halli, and S. König. 2021. Genotype by heat stress interactions for production and functional traits in dairy cows from an across-generation perspective. Journal of Dairy Science 104(9):10029–10039.

Kolmodin, R., E. Strandberg, P. Madsen, J. Jensen, and H. Jorjani. 2002. Genotype by environment interaction in Nordic dairy cattle studied using reaction norms. Acta Agriculturae Scandinavica, Section A-Animal Science 52(1):11–24.

Li, G., S. Chen, J. Chen, D. Peng, and X. Gu. 2020. Predicting rectal temperature and respiration rate responses in lactating dairy cows exposed to heat stress. Journal of Dairy Science 103(6):5466–5484.

Littlejohn, M. D., K. M. Henty, K. Tiplady, T. Johnson, C. Harland, T. Lopdell, R. G. Sherlock, W. Li, S. D. Lukefahr, and B. C. Shanks. 2014. Functionally reciprocal mutations of the prolactin signalling pathway define hairy and slick cattle. Nature communications 5(1):5861.

Marino, L., and K. Allen. 2017. The psychology of cows. Animal Behavior and Cognition 4(4):474–498.

Mattalia, S., A. Vinet, M. P. Calus, H. A. Mulder, M. J. Carabaño, C. Diaz, M. Ramon, S. Aguerre, J. Promp, R. Vallée, B. C. Cuyabano, D. Boichard, E. Pailhoux, and J. Vandenplas. 2023. RUMIGEN: new breeding tools in a context of climate change. Wageningen Academic Publishers.

McLaren, W., L. Gil, S. E. Hunt, H. S. Riat, G. R. S. Ritchie, A. Thormann, P. Flicek, and F. Cunningham. 2016. The ensembl variant effect predictor. Genome Biology 17:1–14.

McManus, C. M., D. A. Faria, C. M. Lucci, H. Louvandini, S. A. Pereira, and S. R. Paiva. 2020. Heat stress effects on sheep: Are hair sheep more heat resistant? Theriogenology 155:157–167.

Meuwissen, T. H. E., B. J. Hayes, and M. E. Goddard. 2001. Prediction of total genetic value using genome-wide dense marker maps. Genetics 157(4):1819–1829. 10.1093/genetics/157.4.1819.

Michael, P., C. R. de Cruz, N. Mohd Nor, S. Jamli, and Y. M. Goh. 2021. The potential of using temperate– tropical crossbreds and agricultural by-products, associated with heat stress management for dairy production in the tropics: a review. Animals 12(1):1.

Misztal, I., L. F. Brito, and D. Lourenco. 2024. Breeding for improved heat tolerance in dairy cattle: methods, challenges, and progress. JDS Communications.

Nadaraya, E. A. 1964. On estimating regression. Theory of Probability & Its Applications 9(1):141–142.

National Research Council. 1971. NRC (1971): A Guide to Environmental Research on Animals. National Academia of science. Washington, DC.

National Research Council (US). 1988. Designing foods: Animal product options in the marketplace. National Academies Press (US).

Nguyen, T. T. T., P. J. Bowman, M. Haile-Mariam, J. E. Pryce, and B. J. Hayes. 2016. Genomic selection for tolerance to heat stress in Australian dairy cattle. Journal of Dairy Science 99(4):2849–2862.

Ojo, T. O., J. Vandenplas, H. A. Mulder, M. L. van Pelt, and M. P. L. Calus. 2024. Genetic Analysis of the Impact of Heat Stress on Fertility Traits in Dairy Cows in the Netherlands. Journal of Dairy Science.

Pook, T., M. Mayer, J. Geibel, S. Weigend, D. Cavero, C. C. Schoen, and H. Simianer. 2020. Improving imputation quality in BEAGLE for crop and livestock data. G3: Genes, Genomes, Genetics 10(1):177– 188.

Pook, T., M. Schlather, G. de los Campos, M. Mayer, C. C. Schoen, and H. Simianer. 2019. HaploBlocker: Creation of subgroup specific haplotype blocks and libraries. Genetics:1045–1061. 10.1534/genetics.119.302283.

Poppe, M., H. A. Mulder, M. L. van Pelt, E. Mullaart, H. Hogeveen, and R. F. Veerkamp. 2022. Development of resilience indicator traits based on daily step count data for dairy cattle breeding. Genetics Selection Evolution 54(1):21.

Pörtner, H.-O., D. C. Roberts, E. S. Poloczanska, K. Mintenbeck, M. Tignor, A. Alegría, M. Craig, S. Langsdorf, S. Löschke, and V. Möller. 2022. IPCC, 2022: Summary for policymakers. 92916915.

Prakapenka, D., Z. Liang, H. B. Zaabza, P. M. VanRaden, C. P. van Tassell, and Y. Da. 2024. A million-cow validation of a chromosome 14 region interacting with all chromosomes for fat percentage in US Holstein cows. International journal of molecular sciences 25(1):674.

R Core Team. 2017. R: A Language and Environment for Statistical Computing, Vienna, Austria. R Foundation for Statistical Computing. https://www.R-project.org/.

Ravagnolo, O., and I. Misztal. 2000. Genetic component of heat stress in dairy cattle, parameter estimation. Journal of Dairy Science 83(9):2126–2130.

Ríus, A. G. 2019. Invited Review: Adaptations of protein and amino acid metabolism to heat stress in dairy cows and other livestock species. Applied Animal Science 35(1):39–48.

Ross, J. W., B. J. Hale, N. K. Gabler, R. P. Rhoads, af Keating, and L. H. Baumgard. 2015. Physiological consequences of heat stress in pigs. Animal Production Science 55(12):1381–1390.

Schaeffer, L. R. 1994. Multiple-country comparison of dairy sires. Journal of Dairy Science 77(9):2671–2678.

Sigdel, A., R. Abdollahi-Arpanahi, I. Aguilar, and F. Peñagaricano. 2019. Whole genome mapping reveals novel genes and pathways involved in milk production under heat stress in US Holstein cows. Frontiers in genetics 10:928.

Simianer, H., T. Pook, and M. Schlather. 2018. Turning the PAGE - the potential of genome editing in breeding for complex traits revisited. World Congress on Genetics Applied to Livestock:190.

Soyeurt, H., P. Dardenne, F. Dehareng, G. Lognay, D. Veselko, M. Marlier, C. Bertozzi, P. Mayeres, and N. Gengler. 2006. Estimating fatty acid content in cow milk using mid-infrared spectrometry. Journal of Dairy Science 89(9):3690–3695.

Soyeurt, H., A. Gillon, S. Vanderick, P. Mayeres, C. Bertozzi, and N. Gengler. 2007. Estimation of heritability and genetic correlations for the major fatty acids in bovine milk. Journal of Dairy Science 90(9):4435– 4442.

St-Pierre, N. R., B. Cobanov, and G. Schnitkey. 2003. Economic losses from heat stress by US livestock industries. Journal of Dairy Science 86:E52–E77.

Su, G., P. Madsen, M. S. Lund, D. Sorensen, I. R. Korsgaard, and J. Jensen. 2006. Bayesian analysis of the linear reaction norm model with unknown covariates. Journal of animal science 84(7):1651–1657.

van Rossum, B.-J., W. Kruijer, F. van Eeuwijk, M. Boer, M. Malosetti, D. Bustos-Korts, E. Millet, J. Paulo, M. Verouden, and R. Wehrens. 2020. Package ‘statgenGWAS’. R package version 1(7).

Vinet, A., S. Mattalia, R. Vallée, C. Bertrand, B. C. D. Cuyabano, and D. Boichard. 2023. Estimation of genotype by temperature-humidity index interactions on milk production and udder health traits in Montbeliarde cows. Genetics Selection Evolution 55(1):4.

West, J. W. 2003. Effects of heat-stress on production in dairy cattle. Journal of Dairy Science 86(6):2131– 2144.

Willer, C. J., Y. Li, and G. R. Abecasis. 2010. METAL: fast and efficient meta-analysis of genomewide association scans. Bioinformatics 26(17):2190–2191.

Yan, G., K. Liu, Z. Hao, Z. Shi, and H. Li. 2021. The effects of cow-related factors on rectal temperature, respiration rate, and temperature-humidity index thresholds for lactating cows exposed to heat stress. Journal of Thermal Biology 100:103041.

